# Systematic Validation of AlphaFold-Predicted Interactomes with LUCIA

**DOI:** 10.64898/2026.04.02.716051

**Authors:** Tianyu Zhang, Julian Kraft, Timothy K. Soh, Malte Kansy, Ingvar Jonsson, Jens B. Bosse

## Abstract

Mapping protein-protein interactions (PPIs) at structural resolution is essential for understanding cellular machinery. While tools like AlphaFold enable proteome-wide structural predictions, the field lacks high-throughput experimental methods to verify these predictions at scale and establish empirical thresholds for confidence metrics, such as the widely used interface predicted Template Modeling (ipTM) score.

We developed LUCIA (LUminescent Cell-free Interaction Assay), a rapid biochemical screening platform bypassing traditional cloning and protein expression to validate direct binary interactions within days. Applying this to herpesviruses, clinically relevant human pathogens with large proteomes, we generated an exhaustive computational interactome of 23,215 AlphaFold-predicted dimer models across three species (HSV-1, HCMV, and KSHV), accessible via our HerpesPPIs database.

Using the HSV-1 interactome as a benchmark, testing 83 high-confidence predicted dimers with LUCIA yielded 23 novel, experimentally validated interactions. By calibrating AlphaFold metrics against this direct binding data, we demonstrated that an ipTM score ≥ 0.80 identifies *bona fide* interactions with a positive predictive value of 77%.

To demonstrate the functional power of this pipeline, we characterized a previously unknown interaction between HSV-1 UL42 and UL8, linking the viral DNA polymerase and helicase-primase complexes. Structure-guided mutations at the predicted and LUCIA-verified UL42-UL8 interface strongly reduced binding *in vitro* and completely abolished viral replication in cell culture.

Combining the scalable LUCIA platform with computational predictions enables researchers to rapidly translate atomic-level models into validated biological mechanisms without the need to solve experimental structures.

## Introduction

Proteins are the molecular drivers of cellular life, yet they rarely function in isolation. From signal transduction to viral replication, biological processes are orchestrated by stable complexes and dynamic macromolecular machines^1,2^. Consequently, understanding cellular machinery requires moving beyond monomeric structures to map the “interactome”, the complete set of physical protein-protein interactions (PPIs). While decades of structural biology have elucidated thousands of individual structures, the experimental determination of protein complexes remains labor-intensive and low-throughput. This bottleneck has left substantial gaps in structural coverage^3,4^, leaving the vast majority of potential protein interfaces unmapped and functionally uncharacterized.

The advent of deep learning-based structure prediction has fundamentally altered this landscape. Models such as AlphaFold (AF)^5,6^ and RosettaFold^7^ demonstrated that atomic-level models can be generated from sequence alone with accuracy rivaling experimental methods, dramatically shrinking the “dark proteome”^5,8^ and enabling structural systems biology^9^. Crucially, these tools extend their capability to protein-protein complexes, enabling proteome-wide screening that predicts not only binding partners but also the precise residues forming the interface^10,11^. This predicted atomic-level detail is transformative: it allows for rational mutagenesis and functional dissection of interactions without the prerequisite of solving experimental structures, a capability previously unattainable at scale.

However, a critical gap persists between *in silico* prediction and experimental verification. Structure prediction tools assign confidence metrics, most notably the interface predicted Template Modeling (ipTM) score, to each modeled complex, and community resources have proposed general confidence tiers to guide interpretation. Yet these thresholds have not been systematically benchmarked against a uniform direct-binding assay, leaving experimentalists without empirically grounded criteria for distinguishing genuine interactions from computational noise. The scale of this challenge is underscored by recent proteome-wide PPI predictions for the human interactome, which catalogue millions of putative complexes^12,13^ but remain largely unvalidated experimentally. Compounding the problem, the experimental methods most commonly used to assess protein interactions are ill-suited to fill this gap. Proximity-based assays such as co-immunoprecipitation, split-luciferase complementation, and NanoBRET report on the spatial proximity of two proteins within a cellular environment, but cannot distinguish direct binary contacts from associations bridged by endogenous cofactors or higher-order complexes, precisely the distinction that validating predicted interfaces demands. Bridging the divide between computational interactomics and biological proof therefore requires a rapid, high-throughput methodology that detects direct physical binding in a cell-free context, using an *E. coli*-derived lysate to minimize interference from host-specific cellular factors.

Developing and benchmarking such a methodology requires a biological system that is computationally tractable, yet complex enough to challenge prediction algorithms and yield biologically meaningful results. Viral proteomes meet all three criteria. With typically tens to low hundreds of proteins, they are orders of magnitude smaller than bacterial or eukaryotic proteomes, making exhaustive pairwise prediction and systematic experimental validation feasible within a single study. At the same time, viruses are far more than convenient model systems: they assemble *bona fide* multi-protein machines, such as polymerase holoenzymes, whose functions can be rigorously tested through reverse genetics. This combination is uniquely powerful because every validated interaction can be directly linked to a functional replication phenotype, connecting structural prediction to biological mechanism in a way that purely computational benchmarks on reference datasets cannot.

We applied this systems-structural biology approach to the *Herpesviridae* family. These large double-stranded DNA viruses establish lifelong infections and cause substantial human disease, including corneal blindness (Herpes Simplex Virus type 1, HSV-1)^14^, congenital neurodevelopmental impairment (Human Cytomegalovirus, HCMV)^15^, and tumorigenesis (Kaposi’s sarcoma-associated herpesvirus, KSHV)^16^. With genomes encoding approximately 80 - 200 proteins depending on the species, herpesviruses are large enough to form complex macromolecular assemblies, such as polymerase and helicase-primase complexes, yet small enough to allow for comprehensive pairwise screening^17,18^. Although essential herpesvirus genes are well-characterized, the interactions involving accessory proteins remain poorly understood, and the complete network of intra-viral PPIs has never been systematically mapped at structural resolution^9,19^. Herpesviruses thus provide an ideal proving ground: small enough for comprehensive analysis, complex enough to challenge prediction algorithms, and medically important such that validated interactions yield actionable insights for future drug development.

Here, we apply this systems-structural approach to three major human-pathogenic herpesviruses. Using AlphaFold-Multimer (AF-M), we predicted every possible intra-viral protein dimer for HSV-1, HCMV, and KSHV, generating 23,215 complex models that are freely accessible through the HerpesPPIs web resource (https://www.herpesfolds.org/herpesppis). To systematically validate these predictions, we developed the LUminescent Cell-free Interaction Assay (LUCIA), a cell-free luminescence-based binding assay that bypasses traditional cloning and purification to test direct binary interactions within days. By benchmarking 83 predicted HSV-1 dimers, we empirically calibrated ipTM score thresholds against direct binding data, defining score ranges that reliably enrich for true interactions versus those that require experimental verification. Crucially, this systematic validation yielded 23 novel protein-protein interactions, significantly expanding the number of known interactions for this virus. Finally, we demonstrate the functional utility of this pipeline by using a predicted interface to design targeted mutations that abolish both binding *in vitro* and viral replication in cell culture, directly linking computational prediction to biological mechanism without recourse to experimental structures. All predictions, validation data, and integrations with published proteomics datasets are available through HerpesPPIs, providing a comprehensive resource for the herpesvirus community and a generalizable blueprint for converting structural interactome maps into mechanistic insight.

## Results

### Proteome-wide *in silico* screen of HSV-1, HCMV, and KSHV dimers identifies conserved interactions

To systematically map herpesvirus interactomes at structural resolution, we used AF-M to predict every possible intra-viral protein dimer for HSV-1 (*Alphaherpesvirinae*), HCMV (*Betaherpesvirinae*), and KSHV (*Gammaherpesvirinae*), yielding 23,215 predicted complex structures in total (Fig. 1A and B; Supplemental Table 1, sheet 1). To make this resource broadly accessible, we expanded our HerpesFolds database with HerpesPPIs, an interactive web platform that allows users to browse complete interactomes, visualize predicted complexes, and inspect confidence metrics together with interface residue details (Fig. 1C). The HerpesPPIs-generated network comprising all PPIs with an ipTM > 0.6 identified conserved interactions across all three herpesvirus subfamilies, underlining AF’s ability to accurately recapitulate the known herpesvirus interactome and underscoring its utility as a powerful tool for prediction-guided hypothesis generation (Fig. 1D). HerpesPPIs further integrates these structural predictions with published experimental datasets, including affinity purification mass spectrometry (AP-MS)^20^, cross-linking mass spectrometry (XL-MS)^21^, and curated PPI repositories (MINT^22^, IntAct^23^, HVInt^24^), enabling users to cross-reference computational predictions with orthogonal evidence and prioritize candidates for functional follow-up (Fig. 1E and F).

**Figure 1:**
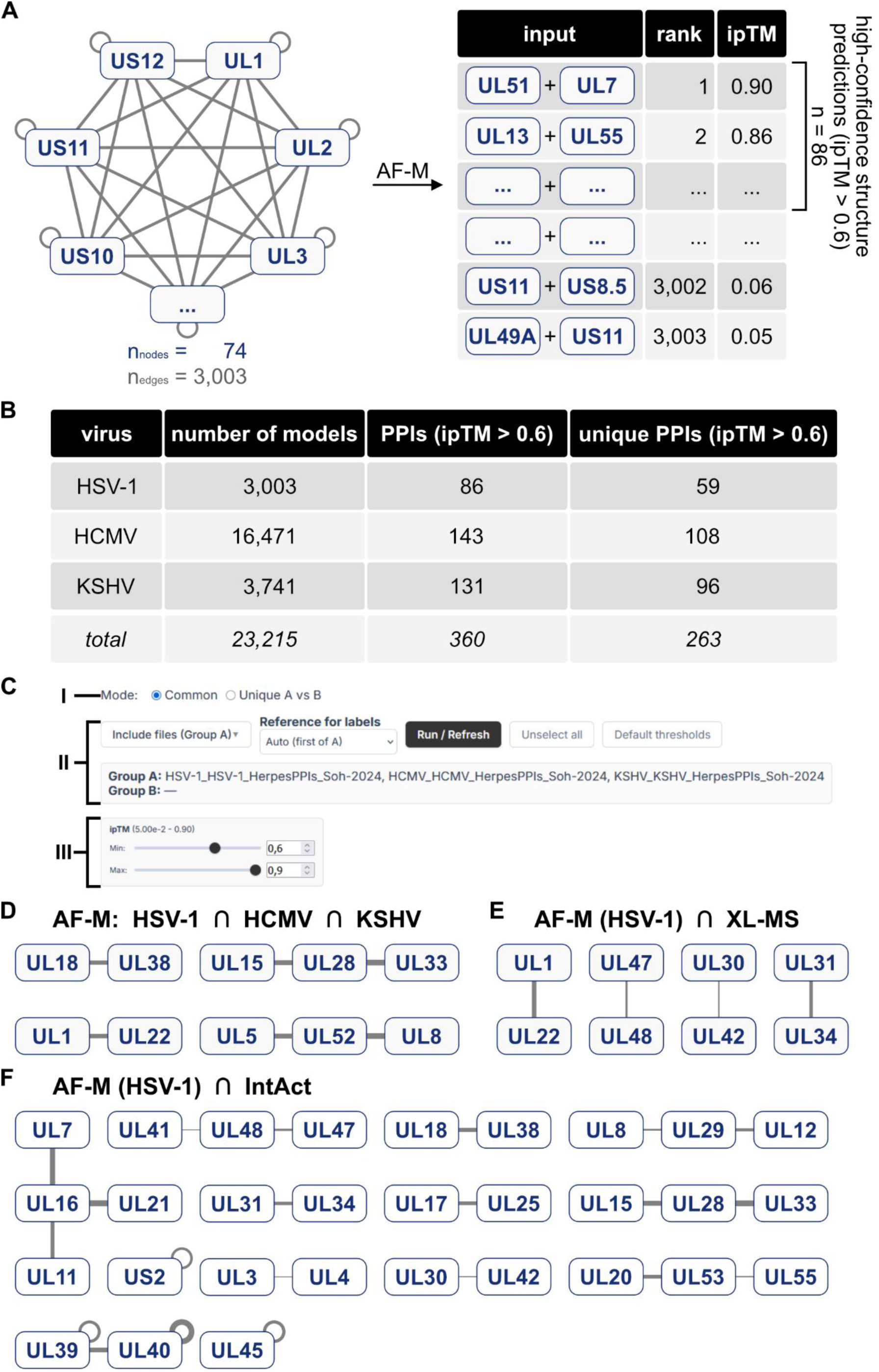
Conservation of predicted interactomes across *Herpesviridae*. **A**, Workflow for *in silico* structure prediction using HSV-1 as an example. Briefly, AF-M was applied to all possible pairwise protein combinations, and the resulting models were ranked according to the AF-M ipTM score. High-confidence models were selected for subsequent *in vitro* validation. **B**, Summary table of the number of pairwise AF-M predictions for HSV-1 (*Alphaherpesvirinae*), HCMV (*Betaherpesvirinae*), KSHV (*Gammaherpesvirinae*), and all viruses combined. For each virus, the total number of AF-M models, the total number of high-confidence PPIs (ipTM > 0.6) as well as the number of species-specific unique high-confidence PPIs (ipTM > 0.6) are provided. **C**, Screenshot of the HerpesPPIs selection board for generating and comparing networks. The interface allows users to define the analysis mode (I), either commonalities or differences in datasets, select the dataset(s) to analyze (II), and define ranges for AF-M and other parameters (III). HerpesPPIs-derived commonality networks depicting high-confidence PPIs (ipTM > 0.6) conserved across all three subfamilies (HSV-1, HCMV, and KSHV) (**D**), or the overlap between AF-M-derived high-confidence HSV-1 PPIs (ipTM > 0.6) and published XL-MS^21^ (**E**) or IntAct databases^23^ (**F**). In all three panels, ∩ means intersection between datasets, nodes represent proteins, and edges denote structural predictions, with edge width proportional to the ipTM score of the corresponding structure prediction.

### LUCIA: A rapid, high-throughput assay for validation of *in silico* predictions

While our computational screen identified hundreds of candidate interactions, translating these predictions into experimentally validated PPIs presents a significant bottleneck. Traditional biochemical approaches to characterize direct PPIs, such as surface plasmon resonance or isothermal titration calorimetry, require individually cloning, expressing, and purifying proteins, a process that typically takes weeks per protein pair and is impractical at the proteome scale. High-throughput cell-based methods, such as yeast two-hybrid or split-reporter assays, can screen large numbers of interactions but still require substantial labor and resources.

To address this gap, we developed the LUminescent Cell-free Interaction Assay (LUCIA), a rapid screen that detects direct protein-protein binding without the need for living cells or conventional protein purification. The key innovation enabling high-throughput application is eliminating all cell-based steps during the initial cloning of the candidate interactors (proteins A and B) (Fig. 2A): rather than cloning the respective coding sequences (CDSs) into expression vectors and transforming *E. coli*, we amplified them by PCR, assembled them into minimal expression backbones via isothermal HiFi assembly, and generated linear concatemeric copies of the resulting minimal expression plasmids through rolling circle amplification (RCA). Following a quality control step to confirm correct assembly and amplification, these templates were directly fed into a prokaryotic cell-free expression (CFE) system, shortening the cloning and protein synthesis pipeline to 2 - 3 days, compared to the weeks required for traditional cloning and purification. Importantly, apart from a plate-reading luminometer, this workflow requires no genetically modified organisms and relies exclusively on standard molecular biology equipment, making LUCIA validation accessible to any molecular biology laboratory.

**Figure 2:**
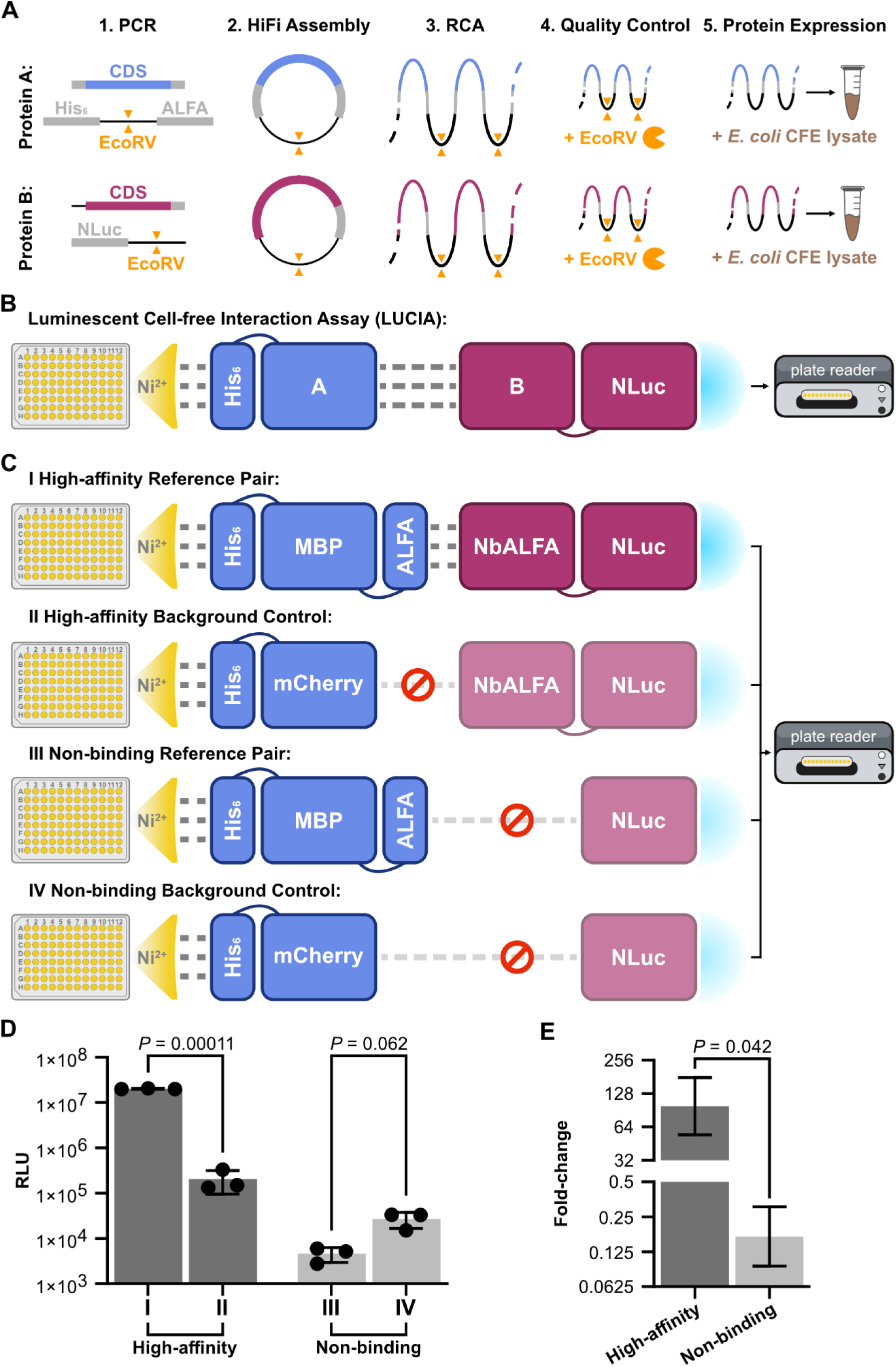
The LUminescent Cell-free Interaction Assay for rapid testing of direct PPIs. **A**, *In vitro* cloning and expression pipeline for candidate interactors (proteins A and B). **B**, Schematic of LUCIA. Protein A is expressed with an N-terminal ALFA-tag and a C-terminal His_6_-tag, and immobilized on a nickel-coated microplate. Protein B is expressed with a C-terminal NLuc and incubated with protein A. Following intermittent washing steps to remove unbound material, retained NLuc-tagged protein B is quantified by bioluminescence using a luminometer, providing a direct readout of PPI. **C**, Schematic of high-affinity and non-binding reference pairs. The high-affinity reference PPI is mediated between the N-terminal ALFA-tag, coupled to His_6_-tagged maltose-binding protein, and a C-terminally NLuc-tagged NbALFA (I), whereas in the non-binding reference pair, maltose-binding protein lacks an ALFA-tag (III). In the respective background controls, the A proteins are replaced with His_6_-tagged mCherry (II and IV). **D**, Bar graph illustrating bioluminescence signals of conditions depicted in (**C**) measured in relative luminescence units (RLUs). Data points of each biological replicate from a distinct sample (*n* = 3) and the mean are shown. Error bars indicate standard deviation. *P* values were calculated with an unpaired one-tailed Welch *t*-test. **E**, Bar graph illustrating FC of bioluminescence signals of high-affinity and non-binding reference pairs relative to their respective background control. Mean FCs are shown. *P* values were calculated with an unpaired one-tailed Welch *t*-test.

In LUCIA, protein A of each candidate pair is expressed with an N-terminal ALFA-tag and a C-terminal His_6_-tag and immobilized on a nickel-coated microplate, after which its putative partner protein B, C-terminally fused to nanoluciferase (NLuc), is added in a subsequent incubation step. Following washing to remove unbound material, retained B protein is quantified by bioluminescence (Fig. 2B). Direct physical interaction between proteins A and B retains NLuc-tagged Protein B on the surface, producing elevated bioluminescence relative to a negative control, in which protein A is replaced with C-terminally His_6_-tagged mCherry, a monomeric fluorescent protein chosen because it is biochemically inert, lacks known binding partners in the tested proteomes, and expresses robustly in prokaryotic cell-free systems. The fold change (FC) of the bioluminescence signal over this mCherry control serves as the primary readout for each candidate interaction.

We validated LUCIA using high-affinity and non-binding reference pairs (Fig. 2C). As a positive reference, we used the biochemically well-characterized interaction between the ALFA-tag and its cognate nanobody NbALFA (Kd = 26 pM)^28^: ALFA-tagged, His_6_-fused maltose-binding protein (ALFA-MBP-His_6_) was immobilized as protein A, and NLuc-fused NbALFA (NbALFA-NLuc) served as protein B. This pair yielded a 98.7-fold increase in signal over the mCherry background control (Fig. 2D and E) consistent with the known picomolar affinity. Notably, AF-M independently predicted this complex with high confidence (ipTM = 0.92). As a non-interacting reference, we paired ALFA-MBP-His_6_ with unfused NLuc (ipTM = 0.19); this combination produced no significant signal above background (Fig. 2D and E), thereby confirming that LUCIA robustly discriminates between interacting and non-interacting pairs and provides a scalable platform for proteome-wide validation.

### Systematic experimental validation reveals ipTM thresholds for high-confidence interactions

Although a growing body of literature has evaluated ipTM and other AF-derived confidence metrics, these analyses are often limited to experimentally predefined sets of candidate interactors or yield inconsistent conclusions^26,27^. This underscores the need for a novel experimental approach that combines sufficient throughput to survey predictions across the full confidence spectrum with the biochemical stringency to confirm direct binary binding, thereby providing system-specific empirical calibration of AF confidence metrics.

Having established LUCIA as a robust platform for detecting direct PPIs, we next applied it systematically to the predicted HSV-1 interactome. To do so, we first selected all 86 predictions with ipTM > 0.6, a threshold commonly cited as the lower boundary for potential interactions^25^, for experimental testing. For each protein pair, the partner exhibiting lower non-specific plate binding was assigned as the NLuc-fused protein B to maximize signal-to-noise ratio (Supplemental Table 2, sheet 1). For transmembrane proteins, soluble ectodomains were expressed where possible to improve compatibility with the CFE system. Of the 86 selected pairs, three could not be tested: two UL36-associated interactions failed repeatedly at the cloning stage, likely due to the large size of the UL36 CDS (∼9.5 kb), and RS1 could not be expressed at detectable levels in the CFE system. The remaining 83 high-confidence predictions were assayed by LUCIA alongside 12 low-confidence predictions (ipTM < 0.6) included as negative controls (Fig. 3A; Supplemental Fig. 1; Supplemental Table 2, sheet 2; Supplemental Table 1, sheet 2).

**Figure 3:**
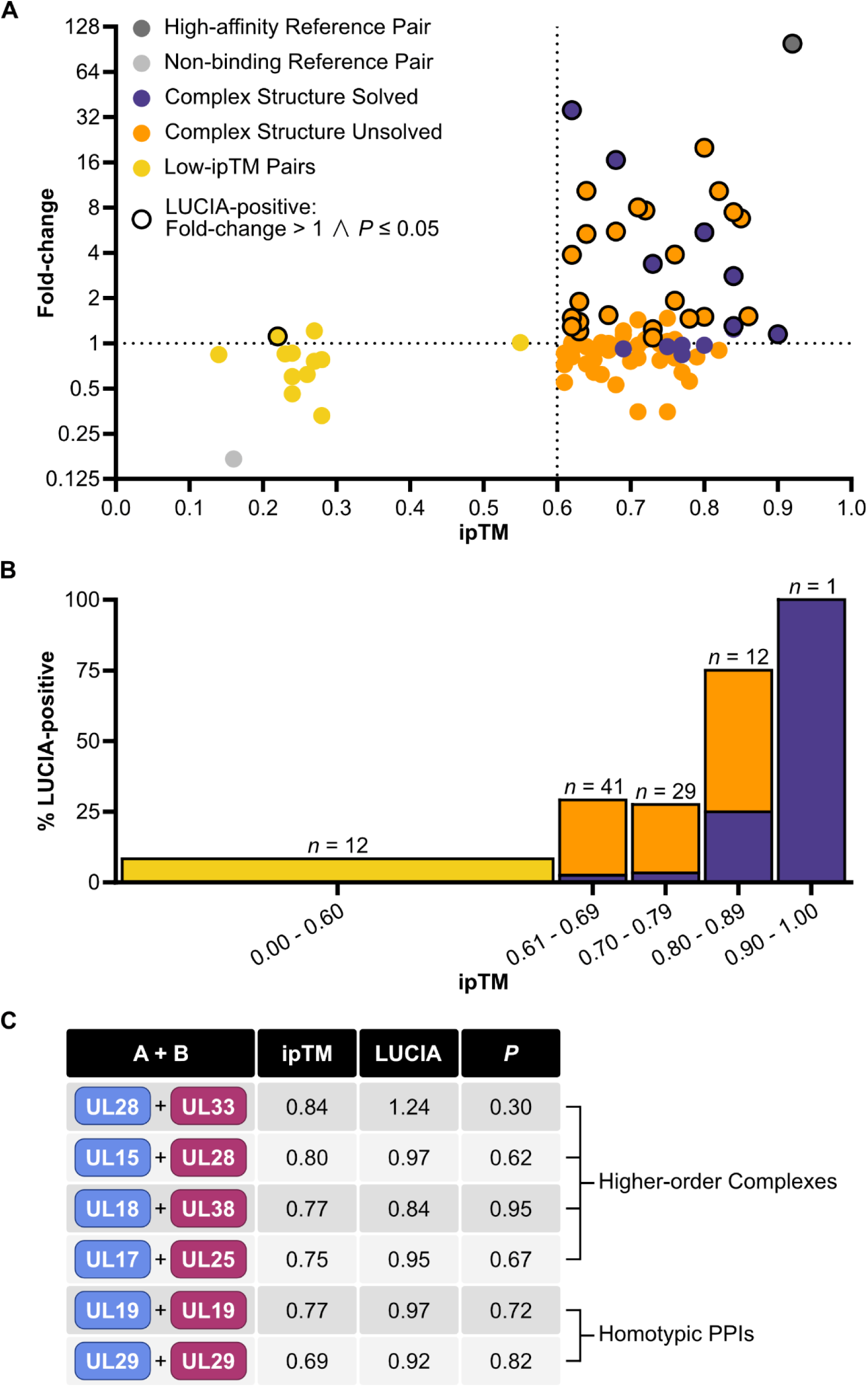
Systematic *in vitro* validation of high-confidence HSV-1 PPIs by LUCIA reveals correlation with ipTM score. **A**, Scatter plot of FC as a function of ipTM score. Shown are 83 of 86 high-confidence PPIs (ipTM > 0.6), including PPIs with (purple circles) and without (orange circles) experimentally solved complex structures. In addition, 12 low-confidence PPIs (ipTM < 0.6; yellow circles), and the high-affinity and non-binding reference pairs (dark and light grey circles, respectively) are displayed. LUCIA-positive PPIs (FC > 1 and *P* ≤ 0.05) are indicated by an additional black outline. Data points represent mean FC values. For each PPI and background control, bioluminescence measurements were performed in three biological replicates from distinct samples, except for UL12-UL13, UL22-US6, UL40-UL40, and UL47-RS1, for which only two replicates were obtained due to low protein yields in the CFE system. **B**, Bar graph showing the fraction of LUCIA-positive HSV-1 PPIs across defined ipTM score intervals. Absolute numbers of PPIs per interval (n) are indicated above each bar. Bar segments are color-coded to indicate experimentally solved (purple), unvalidated (orange), and low-confidence (yellow) interactions. **C**, Summary table of protein pairs with experimentally solved complex structures that were not detected as positive in LUCIA.

The results revealed a clear correlation between ipTM score and experimental validation rate, defining a confidence gradient rather than a single binary threshold. Predictions with ipTM ≥ 0.80 were validated with a positive predictive value (PPV) (*i.e.* number of LUCIA positive/number of ipTM ≥ 0.80) of 77%, whereas those in the 0.61 - 0.79 range showed a substantially lower validation rate of 29%, demonstrating how the false discovery rate scales with ipTM (Fig. 3B). Among the 12 low-confidence controls, only a single pair (8%) produced a positive LUCIA signal, confirming assay specificity. These data establish that ipTM ≥ 0.80 strongly enriches for *bona fide* direct interactions, while the 0.61 - 0.79 interval represents a “grey zone” in which roughly one-third of predictions correspond to genuine binding partners and the remainder require independent experimental verification. We compared our experimental approach with a computational approach using Receiver Operating Characteristic (ROC) analysis on AF structure predictions of a curated set of *E. coli* interacting and non-interacting protein pairs^29^ (Supplemental Table 1, sheet 2). ROC analysis resulted in a threshold of ipTM > 0.47 (Supplemental Fig. 2). This is much lower than our experimental findings, suggesting that a lower cutoff may be appropriate for broad computational discovery, but a substantially higher confidence is needed to reliably predict direct physical binding with a low false discovery rate.

To assess whether recently proposed alternative metrics could improve prioritization within the AF dataset, we evaluated actifpTM^30^ and ipSAE^31^ scores on the same 83 predictions of ipTM > 0.6. Among LUCIA-validated interactions, actifpTM achieved a recall (*i.e.* number of positive scores/number of LUCIA positive) of 80% and ipSAE reached 73% (Supplemental Table 1, sheet 3), suggesting that these metrics capture partially orthogonal features of interface quality and may complement ipTM for candidate ranking in future screens.

To further benchmark LUCIA against structural ground truths, we examined predictions for experimentally solved HSV-1 complexes. For well-behaved dimeric complexes, excluding those known to require a large protein complex, form homo-dimers, or have large disordered domains, LUCIA achieved a recall of 100% (6/6; Fig. 4; Supplemental Table 1, sheet 4). The undetected complexes are informative rather than problematic: they include components of the terminase (UL15-UL28-UL33), triplex (UL18-UL38), and capsid vertex-capping (UL17-UL25) assemblies, all of which require three or more subunits for stable complex formation *in vitro* (Fig. 3C). Their absence from LUCIA is therefore consistent with the known assembly biology of these molecular machines and confirms that the assay reports specifically on binary interfaces rather than higher-order assembly intermediates.

**Figure 4:**
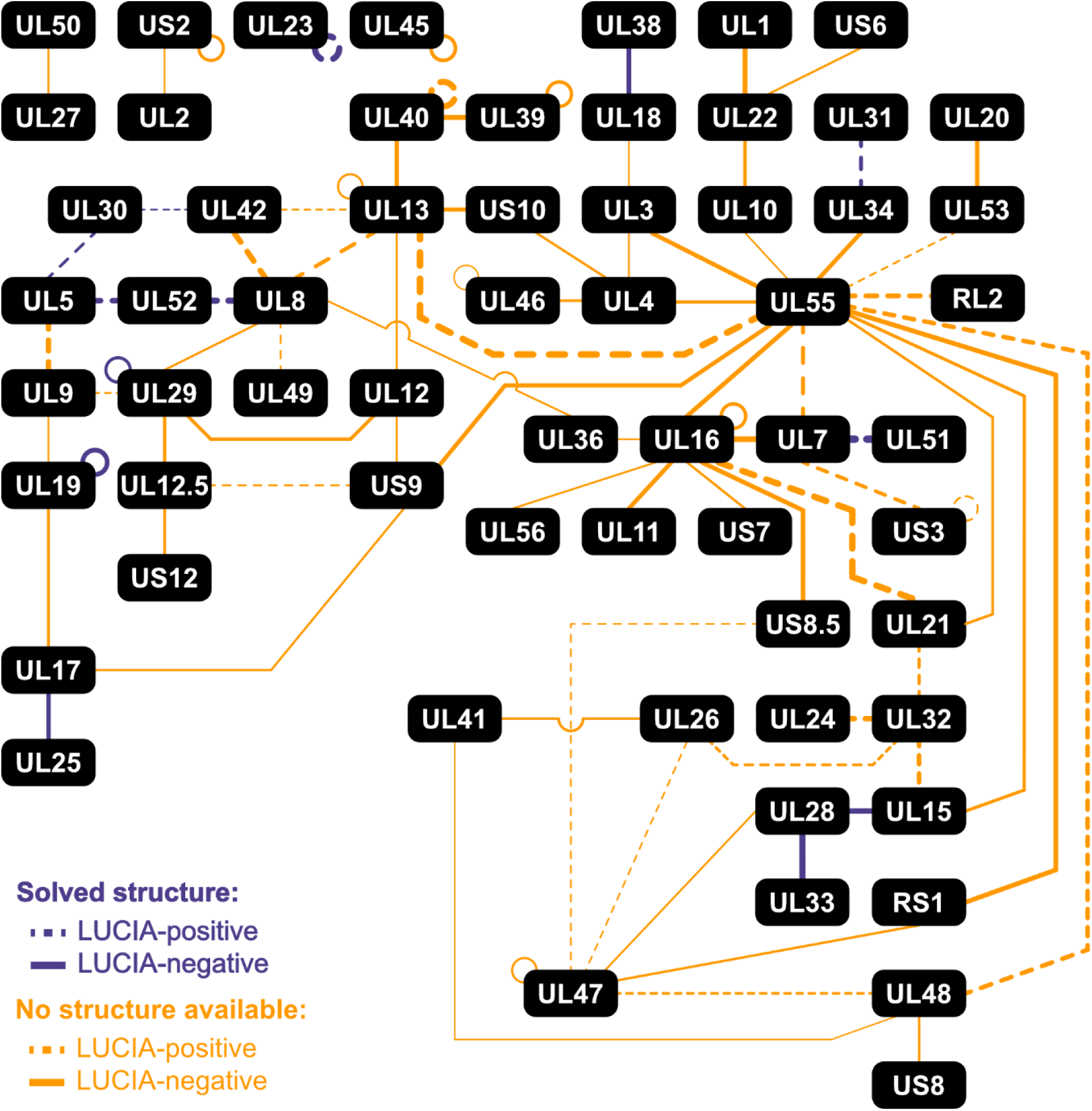
Overlay of AF-M-predicted HSV-1 high-confidence PPI network with LUCIA validation. The 83 high-confidence HSV-1 PPIs (ipTM > 0.6) tested in LUCIA are shown as a network, with structure predictions represented by edges and edge width proportional to the ipTM score. Interactions for which experimentally solved complex structures are available are depicted in purple, whereas interactions lacking solved complex structures are shown in orange. LUCIA-positive PPIs (FC > 1 and *P* ≤ 0.05) are indicated by dashed lines, while LUCIA-negative interactions are shown as solid lines.

Two experimentally confirmed homodimers, UL19-UL19 (PDB: 6CGR)^32^ and UL29-UL29 (PDB: 1URJ)^33^, were not detected by LUCIA (Fig. 3C and 4). This likely reflects a design-inherent limitation of the asymmetric tagging scheme: when a protein forms a strong homodimer, His_6_-tagged protein A molecules self-associate on the plate surface, leaving few unoccupied binding sites for the NLuc-tagged protein B copy. This competition between A-A and A-B binding is expected to specifically suppress signal for symmetric homodimers, and researchers applying LUCIA to other systems should anticipate this constraint when interpreting negative results for predicted self-associations.

We further examined whether high-confidence predictions involving transmembrane proteins reflect genuine extracellular or cytoplasmic interfaces. By comparing full-length and ectodomain-only predictions, we found that complexes with experimentally resolved structures, such as UL34-UL31^34^, maintained high ipTM scores and strong LUCIA signals regardless of construct boundaries, validating the predicted interface. In contrast, other predicted pairs, such as US7-UL16, showed substantially reduced ipTM values upon truncation to soluble domains, and inspection of the full-length models revealed that the predicted binding interface was located entirely within the transmembrane region, a topologically implausible arrangement (Supplemental Fig. 1; Supplemental Table 1, sheet 5; Supplemental Table 2, sheet 3). Transmembrane protein predictions should therefore be interpreted with caution, and we recommend that researchers verify whether the predicted interface maps to a physiologically accessible domain before pursuing experimental follow-up.

In total, systematic LUCIA screening validated 30 direct binary interactions from the 83 high-confidence HSV-1 predictions, 23 of which (77%) lacked previously solved structures and 14 were completely unknown. By assembling these into an experimentally verified, proteome-wide interaction network (Fig. 4), we gained immediate mechanistic insights. For instance, the origin-binding protein UL9 emerges as a crucial interaction hub; its contacts with both UL5 and UL29 suggest it coordinates the recruitment of replication-associated factors. To facilitate further discovery, this LUCIA-validated network can be interactively explored and overlaid with published orthogonal experimental datasets within our https://herpesfolds.org/herpesppis.php web resource. Ultimately, this network demonstrates that coupling exhaustive computational prediction with rapid biochemical screening can substantially expand the known interactome of a complex pathogen in a single campaign, while simultaneously providing the empirical confidence calibration needed to guide similar efforts in other biological systems.

**Supplemental Figure 1:**
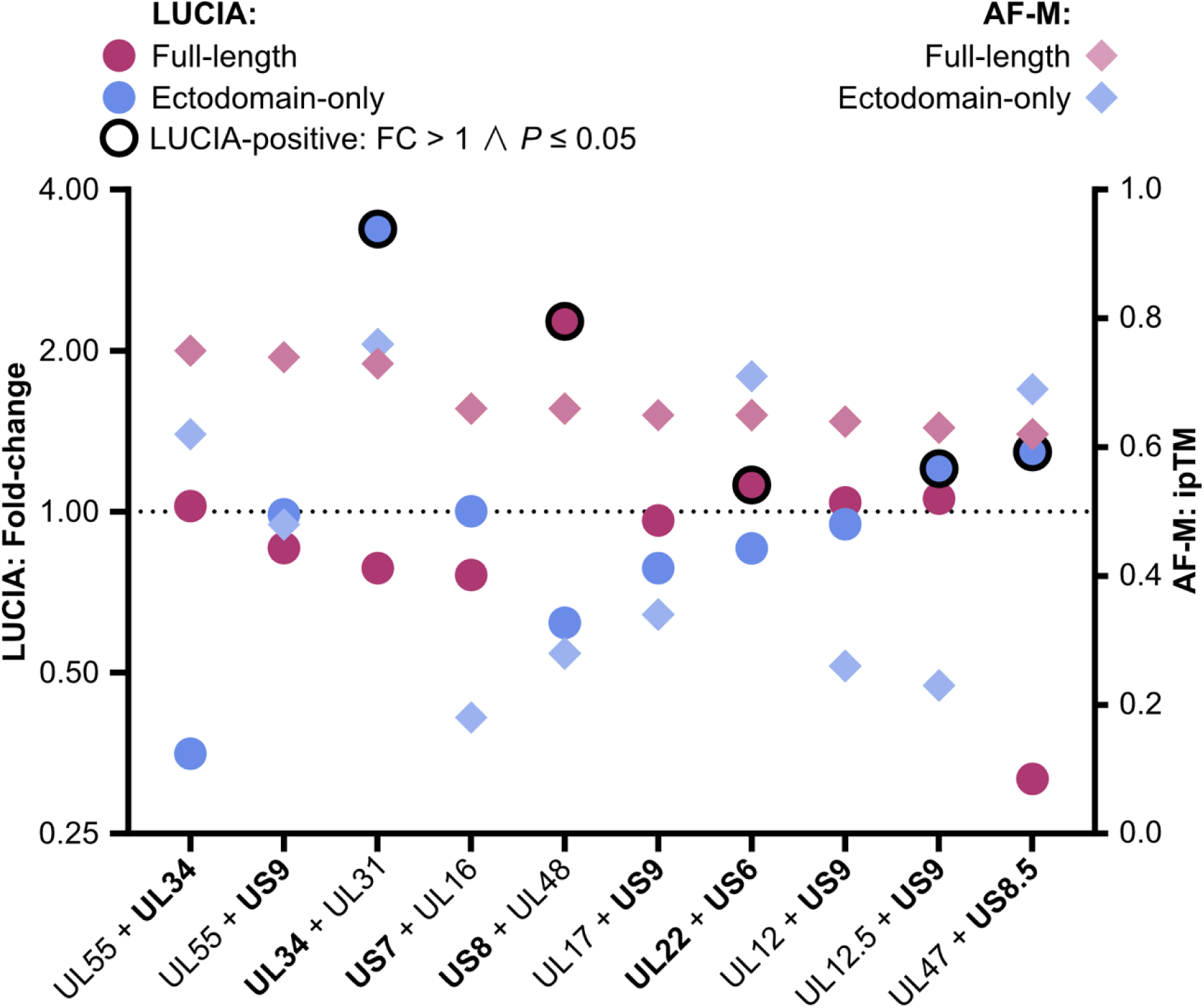
LUCIA and AF-M scores do not correlate for PPIs involving at least one transmembrane protein. Ten high-confidence HSV-1 PPIs containing at least one transmembrane protein (bold) are shown. LUCIA scores (left y-axis) for full-length (purple circles) and ectodomain-only variants (blue circles) are indicated. LUCIA-positive PPIs (FC > 1 and *P* ≤ 0.05) are indicated by an additional black outline. Data points represent mean FC values. For each PPI and background control, bioluminescence measurements were performed in three biological replicates, except for UL22 (ectodomain)-US6 (ectodomain), for which only two biological replicates were obtained due to low protein yields in the CFE system. AF-M scores (right y-axis) for full-length (purple rhombuses) and ectodomain-only variants (blue rhombuses) are indicated.

**Supplemental Figure 2:**
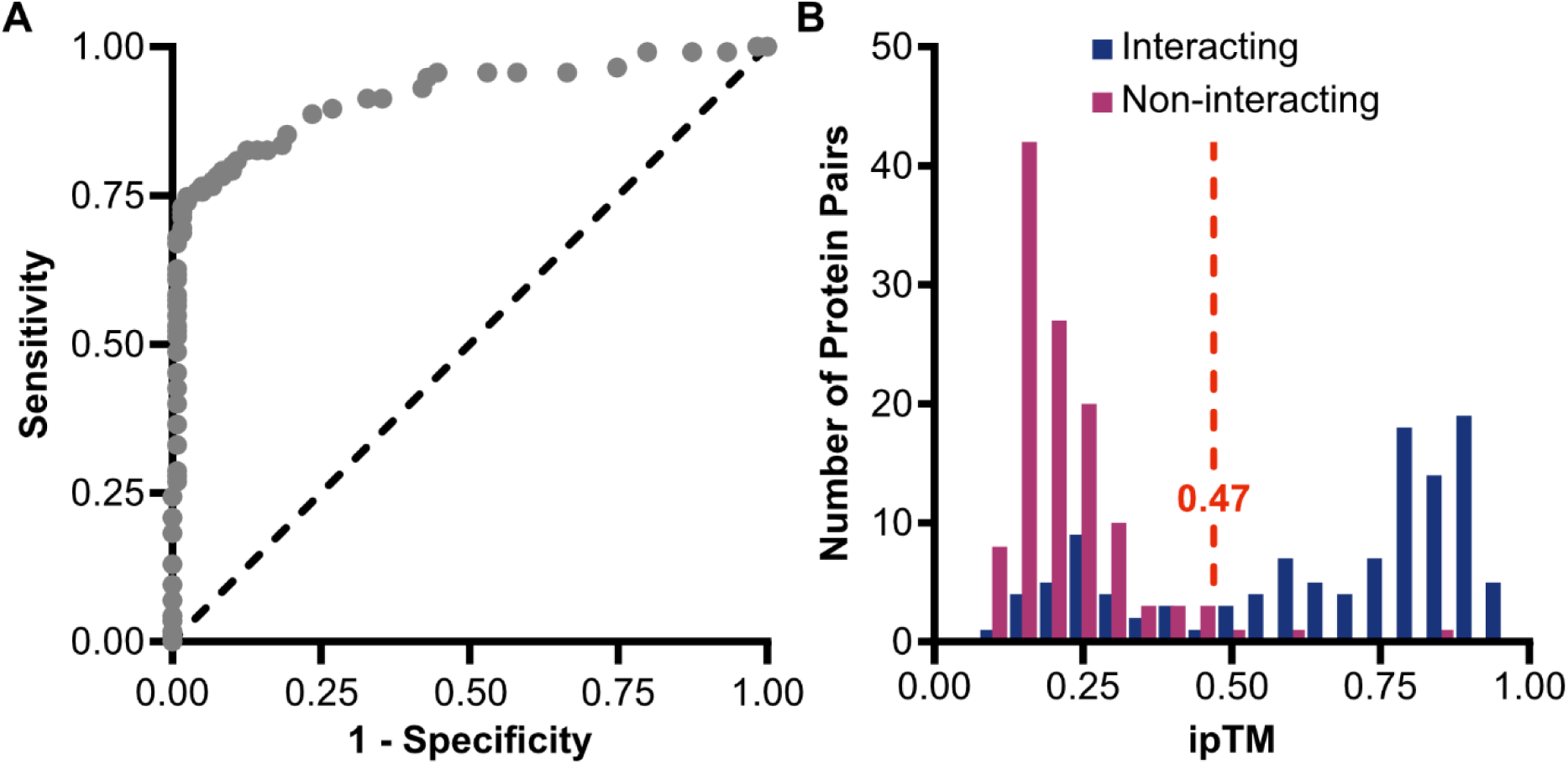
Receiver Operating Characteristic (ROC) analysis of ipTM to identify high-confidence PPIs. **A**, ROC analysis of ipTM on a subset of protein heterodimers from a yeast two-hybrid benchmark database of *E. coli* protein pairs^29^ annotated as interacting or non-interacting. Recall (sensitivity) is plotted against the false positive rate (1 - specificity) across a range of ipTM thresholds. The area under the curve (AUC) is 92%. **B**, Using the identified optimized ipTM threshold of greater than 0.47 (red dotted line), the distributions of the interacting (blue bar graphs) and non-interacting protein pairs (purple bar graphs) relative to the threshold are shown.

### Structure-guided mutagenesis validates predicted interfaces and reveals biological function

To demonstrate that validated predictions yield actionable biological insight, we selected a novel interaction for in-depth functional characterization. The HSV-1 proteins UL42 and UL8 were predicted to form a direct complex with high confidence (ipTM = 0.82; Fig. 5A; Supplemental Fig. 3A-D) despite no prior evidence for a physical interaction. UL42 serves as the processivity factor for the viral DNA polymerase UL30, while UL8 is a core component of the helicase-primase complex^35^. A direct link between these two complexes was identified by cryoEM (PDB: 9YC9)^36^; however, the interaction was mediated by a UL5-UL30 contact. Consistent with this, we detected a UL5-UL30 interaction by AF-M (ipTM = 0.68) and confirmed it by LUCIA (Fig. 4). Notably, the presence of both UL5-UL30 and UL42-UL8 interactions suggests a dual linkage between the polymerase and helicase-primase complexes. Such an arrangement is consistent with the previously proposed trombone model of herpesvirus DNA replication, in which a single helicase-primase complex engages two oppositely oriented polymerase complexes^37^. This architecture would account for the formation of minicircle-like replication intermediates^37^, likely representing looped lagging strand DNA, and provides a mechanistic framework for coupling leading and lagging strand synthesis to replication fork progression (Fig. 5B).

**Figure 5:**
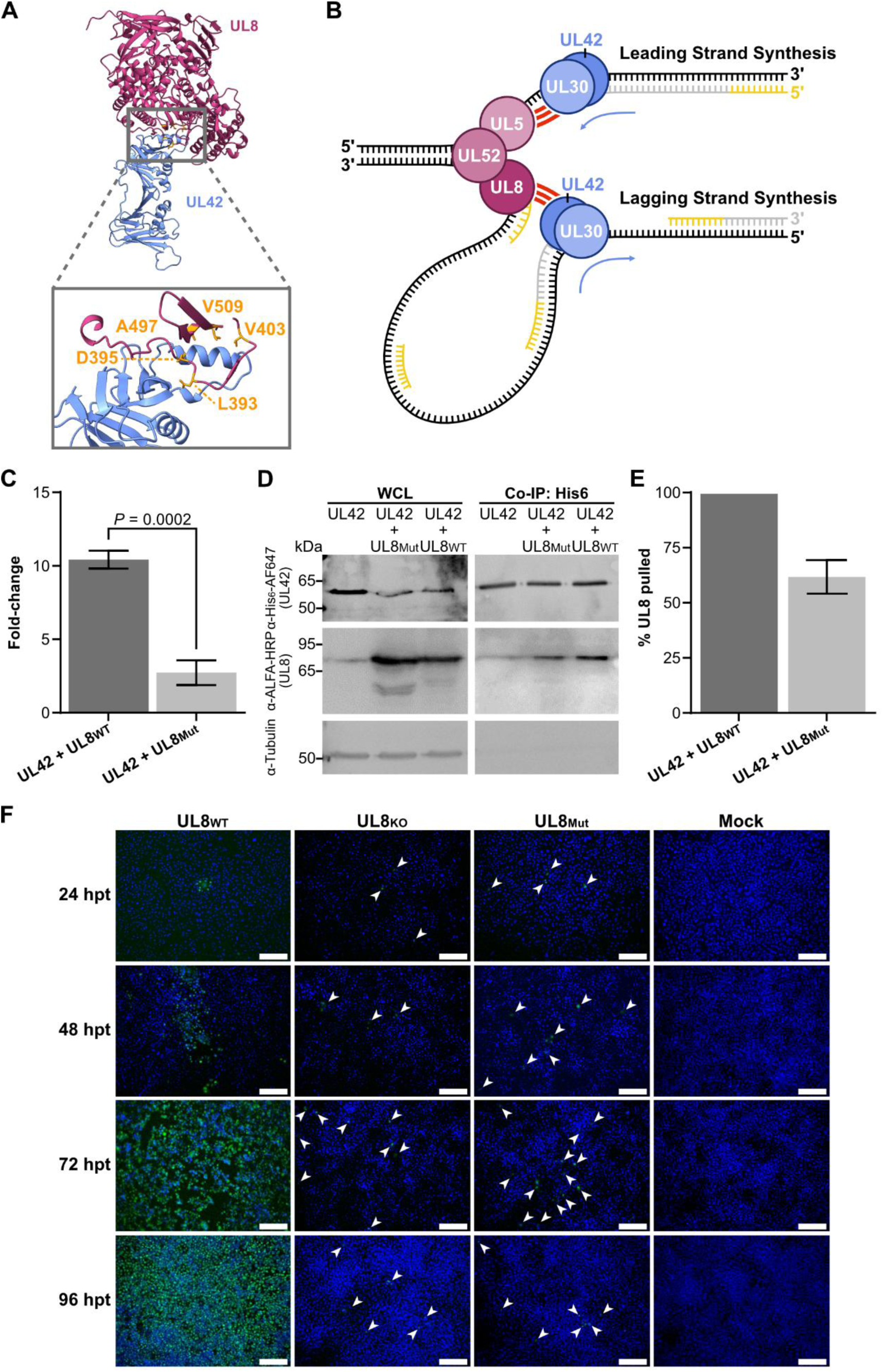
Validation of AF-predicted UL42-UL8 interface. **A**, Structural model of the UL42-UL8 complex predicted by AF-M. UL42 and UL8 are shown in blue and magenta, respectively, with UL8 interface residues highlighted in yellow. Residues 1 - 30 and 318 - 488 of UL42 are omitted for clarity. The entire complex as well as a magnified view of the interface below are shown, indicating key residues contributing to the interaction. The model was visualized in ChimeraX. **B**, Trombone model of HSV-1 DNA replication. A single helicase-primase complex (UL5-UL8-UL52; shades of magenta) is associated with two DNA polymerase complexes in opposing directions (UL30-UL42; shades of blue) via UL5-UL30 and UL42-UL8 interactions (highlighted in red), thereby coupling leading and lagging strand synthesis at the replication fork. Newly synthesized DNA is shown in grey, and RNA primers in yellow. Blue arrows indicate the direction of DNA synthesis by the polymerase complexes. **C**, Bar graph illustrating mean LUCIA FC values of UL42-UL8_WT_ and UL42-UL8_Mut_ combinations. Bioluminescence measurements of each PPI and background control were performed in three biological replicates from distinct samples. Error bars are standard deviation; *P* values were calculated with an unpaired one-tailed Welch *t*-test. **D**, UL42 Co-IP in HEK 293XT cells transiently expressing UL42-His_6_ alone or together with UL8_WT_-ALFA or UL8_Mut_-ALFA. Whole cell lysates (WCL; left) and eluates (right) were analyzed by SDS-PAGE and immunoblotting using the indicated antibodies. **E**, Quantification of UL42 Co-IP from (**D**). Band intensities were determined by grayscale analysis and normalized to the corresponding pulled-down UL42 fractions and UL8 WCL input levels. The pulled UL8_WT_ fraction was set to 100%. Experiments were done with two biological repeats from distinct samples. Error bars are standard deviation. **F**, Vero B4 cells were transfected in triplicate (n = 3) with HSV1(17+)Lox-K33 BACs encoding UL8_WT_, UL8_KO_, or UL8_Mut_, or left untransfected (mock). Cells were fixed at 24 - 96 hours post transfection (hpt), stained for infected cell protein 4 (ICP4; green), and counterstained with Hoechst 33342 to visualize host DNA (blue). Images were acquired using an inverted widefield light microscope. In UL8_KO_ and UL8_Mut_ BAC-transfected cells, white arrowheads indicate punctate ICP4 expression. Representative examples from 1 - 2 recorded fields of view per replicate are shown. Scale bar, 200 μm.

Using the AF-M-predicted UL42-UL8 complex structure, we identified candidate interface residues on the UL8 surface and used FoldX to calculate the predicted destabilization energy (ΔΔG) of individual substitutions (Supplemental Fig. 3E; Supplemental Table 1, sheet 6). We selected five mutations (L393H, D395R, V403R, A497K, V509R) with high predicted ΔΔG values, each introducing charge or steric clashes at the interface. As a computational validation, the quintuple-mutant complex was re-predicted with AF-M, and this yielded a dramatically reduced ipTM of 0.51, consistent with a loss of the interaction at the modeled interface (Supplemental Fig. 3F; Supplemental Table 1, sheet 6). Importantly, the structural integrity of the mutant UL8 was maintained in the predicted complex, with the mutant differing by a root-mean-square deviation of 0.471 Å from the wild-type structure over all 750 residues.

LUCIA confirmed reduced binding experimentally: the wild-type UL42-UL8 pair produced a fold change of 10.31, whereas the quintuple mutant was reduced to 2.62, a ∼75% decrease in signal (Fig. 5C). Notably, the mutant did not abolish binding entirely but retained a low residual signal above baseline. This partial reduction is consistent with specific disruption of the predicted interface rather than global protein misfolding or aggregation, which would be expected to eliminate binding altogether and produce poor protein yields. To orthogonally validate this finding in a cellular context, we performed co-immunoprecipitation (Co-IP) in HEK 293XT cells co-expressing His_6_-tagged UL42 and ALFA-tagged UL8. The five interface mutations reduced co-precipitation by 37.9% compared to wild-type UL8 (Fig. 5D and E). The more moderate reduction observed by co-IP relative to LUCIA is consistent with the distinct biochemical contexts of the two assays: in a cellular environment, endogenous bridging factors or incorporation into larger replication-associated complexes could partially stabilize a weakened binary interface, whereas the cell-free LUCIA format isolates the direct pairwise contact and is therefore more sensitive to interface disruption. Together, both assays independently confirm that the AF-predicted interface residues are critical for UL42-UL8 complex formation, with the graded, rather than binary, loss of binding providing additional confidence that the mutations act through specific interface destabilization.

Finally, we tested whether the UL42-UL8 interaction is functionally required during infection. Using bacterial artificial chromosome (BAC)-based reverse genetics, we generated an HSV-1 mutant carrying the five interface substitutions in UL8, along with a complete UL8 deletion as a control. Upon BAC reconstitution by transfection, neither the interface mutant nor the UL8 knockout produced plaques; only single infected cells were observed even at 96 hours post transfection, whereas cells transfected with wild-type HSV-1 BAC were completely infected (Fig. 5F; Supplementary Fig. 4). The interface mutant thus phenocopied the complete knockout, demonstrating that the UL42-UL8 interaction, specifically the AF-predicted and LUCIA-verified contact surface, is essential for productive viral replication. This data illustrates the power of the presented workflow: From initial computational prediction to a replication-null phenotype in a viral infection model, this workflow required only weeks, illustrating how the combination of structural prediction and rapid cell-free validation using LUCIA can accelerate the pace from hypothesis to functional readout.

**Supplemental Figure 3:**
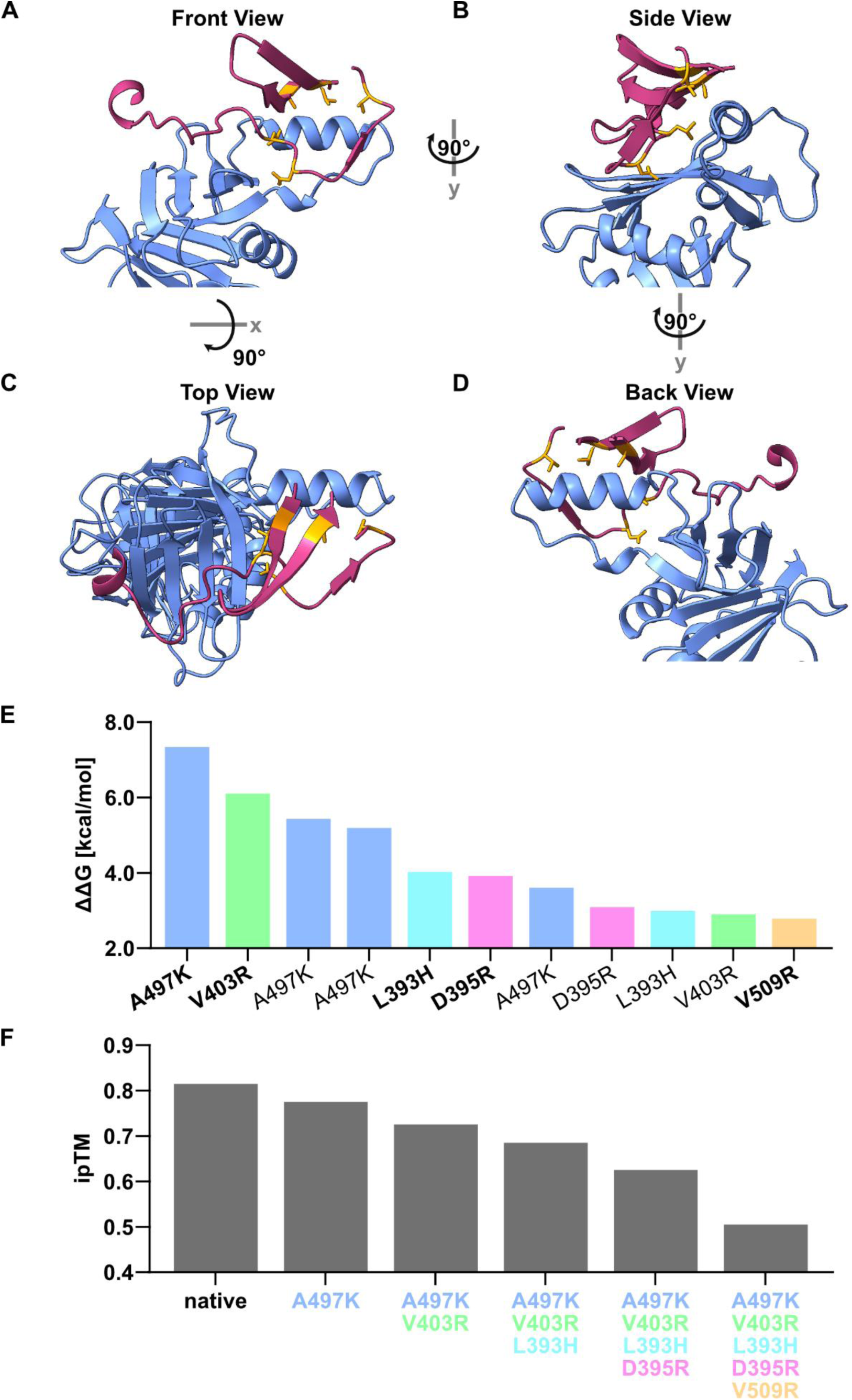
FoldX-based identification of critical UL8 interface residues in the UL42-UL8 complex. **A**-**D**, AF-M-predicted structure of the UL42-UL8 interface shown from front (**A**), side (**B**), top (**C**), and back (**D**) views. UL42 and UL8 are shown in blue and magenta, respectively, with UL8 interface residues highlighted in yellow. Residues 1 - 30 and 318 - 488 of UL42 are omitted for clarity. The model was visualized in ChimeraX. **E**, Bar graph depicting the top 11 most destabilizing UL8 amino acid substitutions within the UL42-UL8 complex, as identified by FoldX. Substitutions were ranked by predicted increase in ΔΔG. UL8 substitutions affecting the same residue are color-coded. The five UL8 substitutions selected for UL8_Mut_ (LUCIA) and HSV1(17+)Lox-K33-UL8_Mut_ (BAC transfection) are indicated in bold. **F**, Bar graph depicting AF-M ipTM scores of the UL42-UL8 wild-type complex (native) and variants carrying sequentially combined destabilizing substitutions.

**Supplemental Figure 4:**
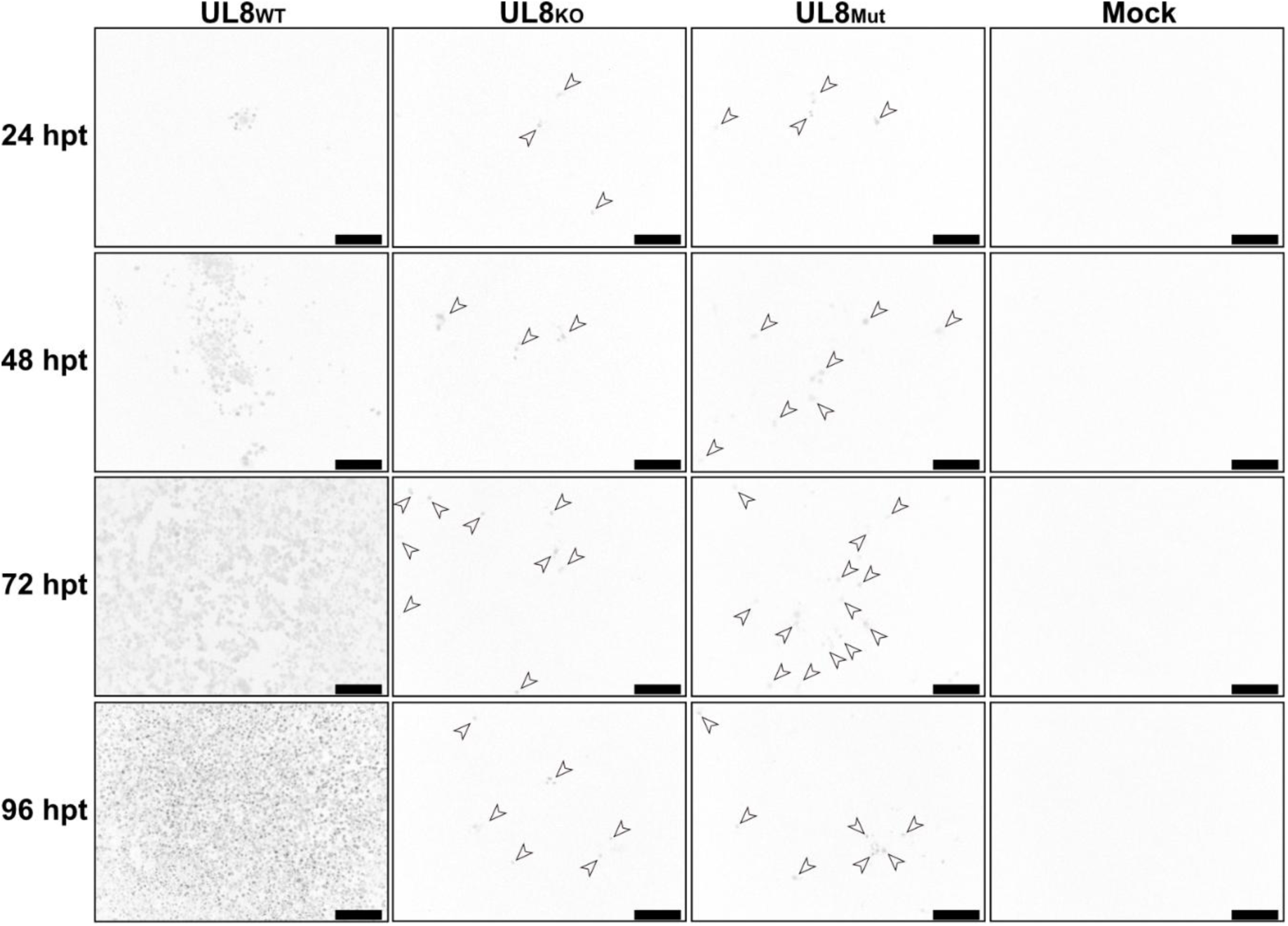
ICP4-positive Vero B4 cells following transfection with HSV1(17+)Lox-K33 BACs encoding UL8_WT_, UL8_KO_, or UL8_Mut_. Vero B4 cells were transfected in triplicate (n = 3) with HSV1(17+)Lox-K33 BACs encoding UL8_WT_, UL8_KO_, or UL8_Mut_ or left untransfected (mock). Cells were fixed at 24 - 96 hpt, stained for ICP4, and counterstained with Hoechst 33342 to visualize all nuclei. Only ICP4 signals are shown, displayed in grayscale and inverted. Images were acquired using an inverted widefield light microscope. In UL8_KO_ and UL8_Mut_ BAC-transfected cells, arrowheads indicate punctate ICP4 expression. Representative examples from 1 - 2 recorded fields of view per replicate are shown. Scale bar, 200 μm.

## Discussion

The past five years have witnessed an explosion in proteome-wide structure prediction^38^, yet the field has lacked systematic approaches to convert these computational resources into experimentally validated interaction networks. We addressed this gap by coupling exhaustive AF-M predictions across the herpesvirus family with LUCIA, a rapid cell-free assay that measures direct binary protein-protein binding. This integrated pipeline moves from DNA sequence to validated interaction in days rather than the months required by conventional cloning and purification workflows. By correlating LUCIA outcomes with AF confidence metrics across a large, uniformly tested set of viral dimers, we established an empirical calibration of the ipTM score against experimental binding data. This systematic evaluation of high-confidence predictions yielded 23 novel protein-protein interactions and an interactome that recapitulated all known conserved interactions, underscoring both the predictive power of AF-M and the sensitivity of LUCIA as a discovery tool. Crucially, we demonstrated that predicted interfaces are not merely computational abstractions: targeted mutagenesis of AF-identified contact residues in the newly identified UL42-UL8 complex strongly reduced binding *in vitro* and abolished productive viral replication in cells, directly linking structure prediction to biological function. The scalability of LUCIA renders this pipeline broadly applicable to proteome-wide interaction studies. Beyond this proof of concept, the interactome itself offers a systems-level view of herpesvirus biology and lays the groundwork for a deeper molecular understanding of a family of clinically relevant human pathogens.

A key practical outcome of this work is the empirical calibration of ipTM confidence tiers that experimentalists can apply when prioritizing AF-predicted interactions in their own systems. Our data indicate that predictions scoring ipTM ≥ 0.80 can be pursued with high confidence for functional follow-up, whereas the 0.61 - 0.79 interval should be treated as a discovery zone in which approximately one-third of predictions correspond to genuine direct interactions and the remainder require independent experimental verification. Below 0.60, our negative-control set, our data suggest that false positive predictions are rare. These empirically grounded ranges formalize what previous studies applied heuristically: Kawaguchi *et al.* used AF to identify interactions among nuage proteins^39^, Homma *et al.* screened pathogen-secreted effectors against plant immune receptors^40^, and Abulude *et al.* combined STRING network analysis with AF-M for predatory bacteria^41^. Each of these studies relied on ad hoc ipTM cutoffs calibrated to their specific biological context. Our systematic benchmarking against a uniform direct-binding assay now provides a quantitative reference that these and future efforts can use to estimate expected validation rates before committing to experimental follow-up.

Beyond methodological benchmarking, our functional characterization of the UL42-UL8 complex represents a biologically significant finding in its own right. The fact that disruption of this single predicted interface phenocopied a complete UL8 knockout suggests that the UL42-UL8 contact is not merely an accessory interaction but a functionally essential connection within the replisome architecture. Together with the additional UL5-UL30 interaction^36^, linking the helicase-primase complex (UL5-UL8-UL52) to the polymerase complex (UL30-UL42) through both computational prediction and experimental validation lends strong support to a trombone model of herpesvirus replication^37^. Whether the UL42-UL8 interface is conserved across herpesviruses, whether it is maintained during active replication, and whether it represents a viable target for antiviral intervention, particularly for decoupling polymerase processivity from fork progression, are important questions for future investigation.

The prediction-driven approach we describe is inherently comprehensive, making structural interactomics a powerful complement to mass spectrometry (MS)-based structural proteomics. While MS has transformed our understanding of herpesvirus architecture, it remains biased toward abundant, stable assemblies. Virion-wide cross-linking MS (XL-MS) of HCMV revealed interaction networks organized around high-copy tegument scaffolds^42^, while in-cell XL-MS has mapped hundreds of HSV-1 virus-host interfaces (SHVIP)^21^. However, these studies consistently show that abundant structural proteins dominate cross-link recovery, systematically underrepresenting interactions involving low-abundance accessory and regulatory proteins^43^. Several of our highest-confidence LUCIA-validated dimers involve precisely such accessory proteins, underscoring the value of prediction-driven discovery for interactions that escape conventional biochemical detection. Conversely, XL-MS captures transient and context-dependent contacts that static dimer predictions may miss. The two approaches are therefore complementary, and overlaying both datasets within HerpesPPIs allows users to identify interactions supported by convergent evidence from orthogonal methods.

Our first-generation implementation deliberately focuses on pairwise dimer predictions, although many herpesvirus proteins function within higher-order assemblies. We view these dimeric predictions as projections of quaternary structure onto individual binary interfaces: useful for identifying contact surfaces and mutational hotspots, but not yet sufficient to reconstruct complete molecular machines. This reflects a current computational constraint, as structural prediction tools lose accuracy with increasing chain number, complex size, and unknown stoichiometries. However, recent work demonstrates that large assemblies can be built up from accurately predicted subcomponents using Monte Carlo tree search and related combinatorial strategies^44^. The dimer-level interactome we report provides a natural scaffold for such hierarchical assembly modeling and already highlights specific interfaces within trimeric complexes and capsid components that merit future higher-order analysis.

As with any experimental platform, LUCIA has inherent limitations that define its domain of applicability. Expression in *E. coli* lysate likely prevents bridging of complexes with host proteins but leads to a lack of eukaryotic post-translational modifications, including glycosylation and phosphorylation, that are critical for the folding or interaction competence of some proteins, potentially generating false negatives for modification-dependent complexes. The asymmetric tagging scheme, while enabling clean discrimination of heterodimeric binding, creates a specific blind spot for homodimers, as demonstrated by UL19 and UL29. Strong self-association of His_6_-tagged protein A on the plate surface can occlude binding sites for the NLuc-tagged copy, suppressing signal even for *bona fide* self-interactions. More broadly, any interaction that depends on three or more subunits for stable assembly, as exemplified by the terminase, triplex, and capsid vertex complexes in our dataset, will be underrepresented in a pairwise assay by design. However, these limitations can be mitigated with due diligence. Resources such as UniProt^45^ and homology-based tools^46,47^ can flag potential confounders and thereby inform which candidates will require alternative validation strategies, such as eukaryotic cell-free expression systems, reconstitution with additional subunits, or cell-based proximity assays.

Looking ahead, this framework is readily extensible in both biological scope and experimental capability. LUCIA can be adapted to test tertiary or higher-order assemblies by combining multiple subunits in the same reaction or repurposed for competition assays in which small molecules or peptides are screened for their ability to disrupt validated interfaces, a format directly relevant to early-stage drug discovery. From a translational perspective, the structurally resolved, replication-linked PPIs uncovered here represent a rich set of candidate antiviral targets. Decades of progress in PPI pharmacology have overturned the view that protein-protein interfaces are intrinsically undruggable, with multiple clinically advanced PPI modulators now in development across therapeutic areas^48–51^. By pinpointing key residues in essential herpesvirus complexes, exemplified by the UL42-UL8 interface whose disruption abolishes replication, our HerpesPPIs database provides rational starting points for the design of small molecules, peptides, or biologics targeting viral protein interfaces. Because the entire LUCIA pipeline relies on openly available prediction tools and standard molecular biology equipment, we expect this framework to be broadly applicable to other viral families, host-pathogen systems, and ultimately any genome for which a comprehensive proteome or interactome map is available.

## Materials and Methods

### Cell Culture and Viruses

Vero B4 (DSMZ ACC 33) and HEK 293XT (Takara) cells were maintained in Dulbecco’s Modified Eagle Medium (DMEM, ThermoFisher, 31966-047) supplemented with 10% fetal bovine serum (FBS Good, PAN-Biotech). All cells were cultured at 37°C in a humidified atmosphere containing 5% CO2. The HSV-1 strain 17+ BAC HSV1(17+)Lox-K33 was kindly provided by Beate Sodeik^52,53^.

### BAC Mutagenesis and Sequencing

The HSV1(17+)Lox-K33 BAC served as the backbone for generating a UL8 CDS knockout mutant (UL8_KO_) via *en passant mutagenesis*^54^. To generate the UL8 interface mutant (UL8_Mut_), the following specific amino acid substitutions were introduced into the UL8 locus of the UL8_KO_ BAC: L393H, D395R, V403R, A497K, V509R. BAC DNA was extracted from bacteria cultures utilizing a plasmid DNA Midiprep Kit (Macherey-Nagel, 740410.100) and subsequently verified by Nanopore sequencing. Primers utilized for BAC mutagenesis are listed in Supplementary Table 2 Sheet 4.

### Protein Sequences

Protein sequences were retrieved from the following UniProt reference proteomes: HSV-1 (strain 17): UP000009294, HCMV (strain Merlin): UP000000938, KSHV (strain GK18): UP000000942.

### Structure Predictions

Protein structure predictions were performed using LocalColabFold v1.5.1^55^. For each protein target, unless otherwise specified, the following parameters were applied: Models: 5 independent models generated per run. Recycling: 3 recycles. Early Stopping: Stop-at-score threshold of 100.

### Determination of ipTM Threshold

To establish a robust ipTM threshold for identifying PPIs, we used a subset of a validated benchmark dataset^29^ comprising of 115 verified interacting protein pairs and 119 non-interacting pairs. The predictive performance of ipTM was evaluated using ROC analysis. To determine the optimal cut-off value, we applied the Youden Index, which maximizes the sum of sensitivity and specificity.

### Protein Expression and Purification from *E. coli* Lysate

For the generation of standard curves and blocking reagents, various recombinant proteins, including N-terminally ALFA-tagged and C-terminally His_6_-tagged maltose-binding protein (ALFA-MBP-His), anti-ALFA nanobody fused to C-terminally tagged NanoLuciferase (NbALFA-NLuc-StrepII), and C-terminally His_6_-tagged mCherry (mCherry-His_6_), were expressed in *E. coli* BL21 using the pET21a(+) vector.

Bacterial cultures were grown in Terrific Broth medium at 37°C to an OD600 of 0.6. Protein expression was subsequently induced with 50 μM filter-sterilized isopropyl β-D-1-thiogalactopyranoside (IPTG; Sigma Aldrich, I6758). Following induction, cultures were incubated at 20°C with shaking at 110 rpm for 16 hours. Cells were harvested by centrifugation and resuspended in NSP12 lysis buffer [20 mM Tris-HCl, 0.1 mM EDTA-NaOH (pH 8.2), 50 mM NaCl, 0.01% (v/v) ZnCl2, 5% (v/v) glycerol] supplemented with 20 mM imidazole and cOmplete Protease Inhibitor Cocktail (Roche, 11836170001). ALFA-MBP-His and mCherry-His_6_ were purified using HisPur Ni-NTA Resin (Thermo Fisher Scientific, 88222), while NbALFA-NLuc-StrepII was purified using Strep-Tactin XT (IBA Lifesciences, 2-5010-025), both according to the manufacturers’ protocols.

Protein purity was verified by SDS-PAGE with Coomassie Brilliant Blue staining and western blot analysis. For immunoblotting, ALFA-MBP-His_6_ was detected using an Alexa Fluor 647-conjugated anti-His_6_ tag antibody (1:5,000; BioLegend, 362611), mCherry-His_6_ was detected using an anti-RFP mouse IgG2c primary antibody (1:2,000; chromotek, 6G6) and a mouse IgG1-AF647 secondary antibody (1:5,000; ThermoScientific, A21236), and NbALFA-NLuc-StrepII was detected using an anti-Strep-tag antibody (1:5,000; GenScript, A01732S) (Supplemental Fig. 5). Chemiluminescent signals were captured using a FUSION FX7 EDGE 18.11 imaging system. Final protein concentrations were determined via spectrophotometry using a NanoDrop spectrophotometer.

**Supplemental Figure 5.**
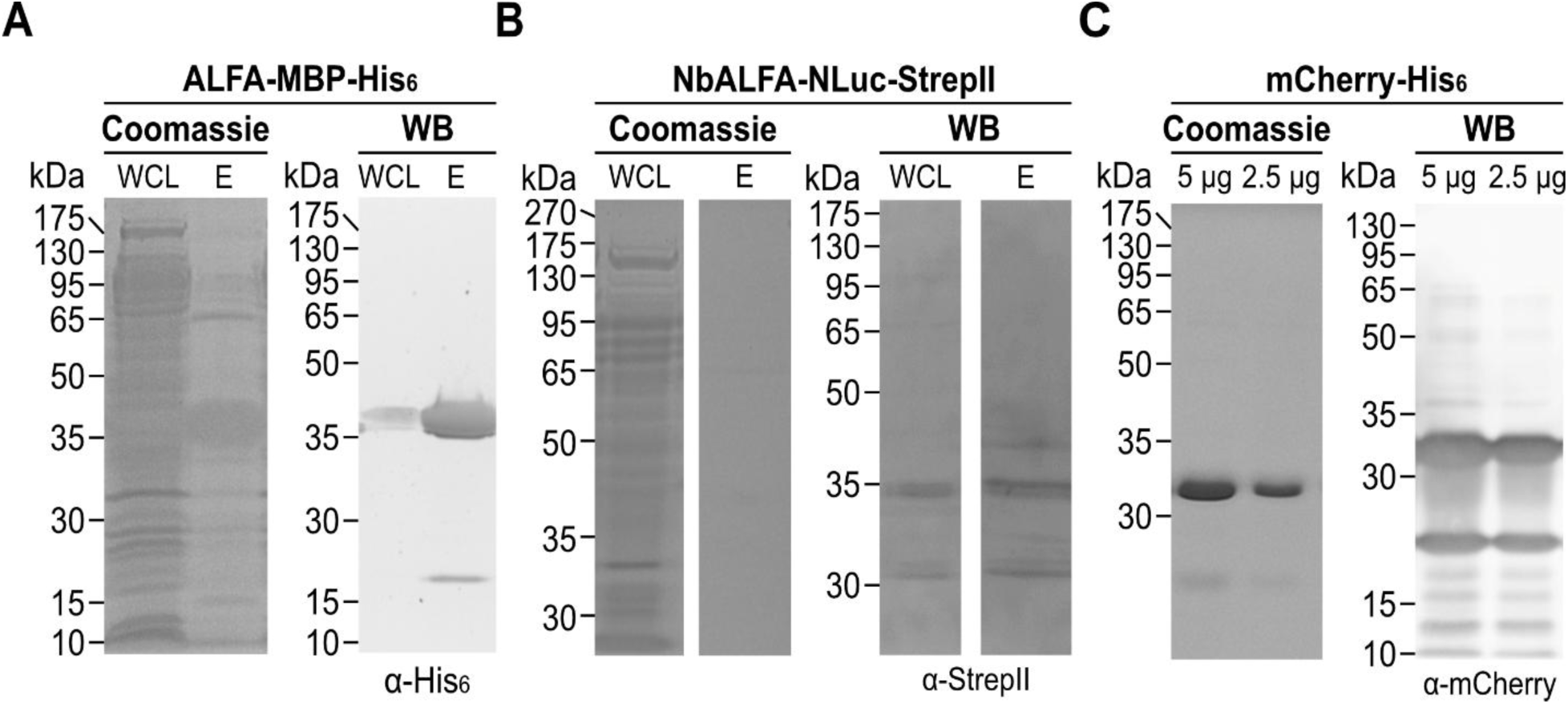
Purification of proteins from *E. coli* lysate. ALFA-MBP-His_6_ (**A**), NbALFA-NLuc-Strep (**B**) and mCherry-His_6_ (**C**) proteins were expressed in *E. Coli* strain BL21. Proteins were purified from bacterial lysate using affinity beads. Purity of the proteins was validated with Coomassie Blue staining (left panel) and western blotting (right panel). WCL, whole cell lysate. E, elution of purified protein product.

### Amplification of HSV-1 CDS and Generation of Linear CFE Templates

HSV-1 CDSs were amplified by PCR using KOD DNA Polymerase (Merck, 71085-3). Primers were designed with specific overhangs to facilitate Gibson-style assembly into universal CFE vector backbones (Supplemental Table 2, sheet 5). The specific overhang sequences used were: A proteins: 5′-GGTAGTAGCTCCTCTGGCGGT-3′ (forward) and 5′-(CC)AGAACCGCCACCGCCTGA-3′ (reverse); B proteins: 5′-(T)TAAGAAGGAGATATACCATG-3′ (forward) and 5′-TCAGGCGGTGGCGGTTCTGG-3′ (reverse). Following amplification, PCR products were resolved by agarose gel electrophoresis to verify size (Supplemental Fig. 6) and subsequently purified using a NucleoSpin Gel and PCR Clean-UP kit (Macherey-Nagel, 740609.250). The purified CDS fragments were inserted into the respective vector backbones via HiFi DNA Assembly (NEB, E2621X) at 50°C for 30 minutes. To generate high-yield DNA templates for CFE, 5 μL of the HiFi assembly product was used as a template for RCA using the phi29-XT RCA Kit (NEB, E1603L) according to the manufacturer’s instructions. The fidelity and yield of the resulting RCA products were confirmed by EcoRV restriction digestion and agarose gel electrophoresis (Supplemental Fig. 7).

**Supplemental Figure 6:**
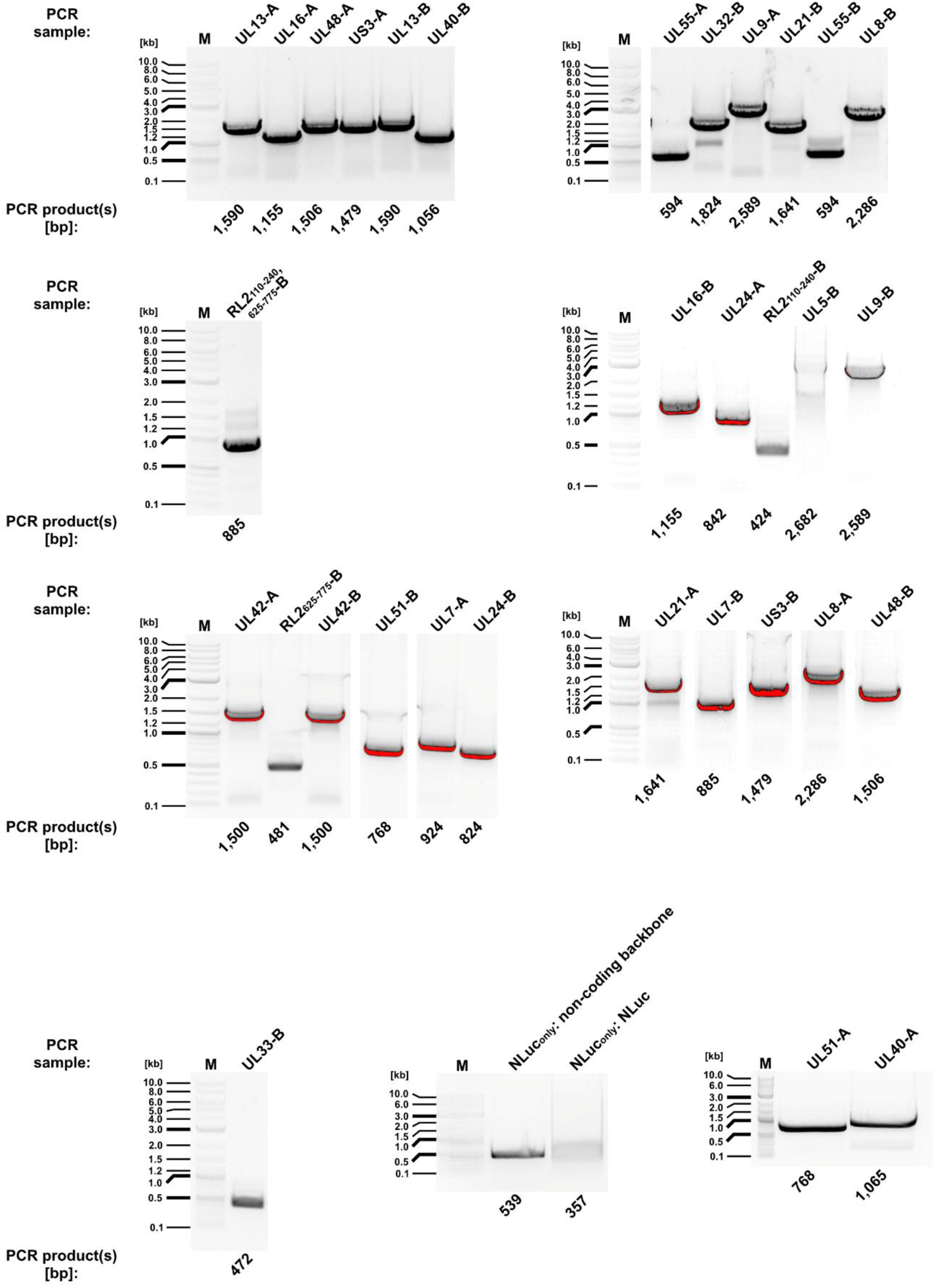
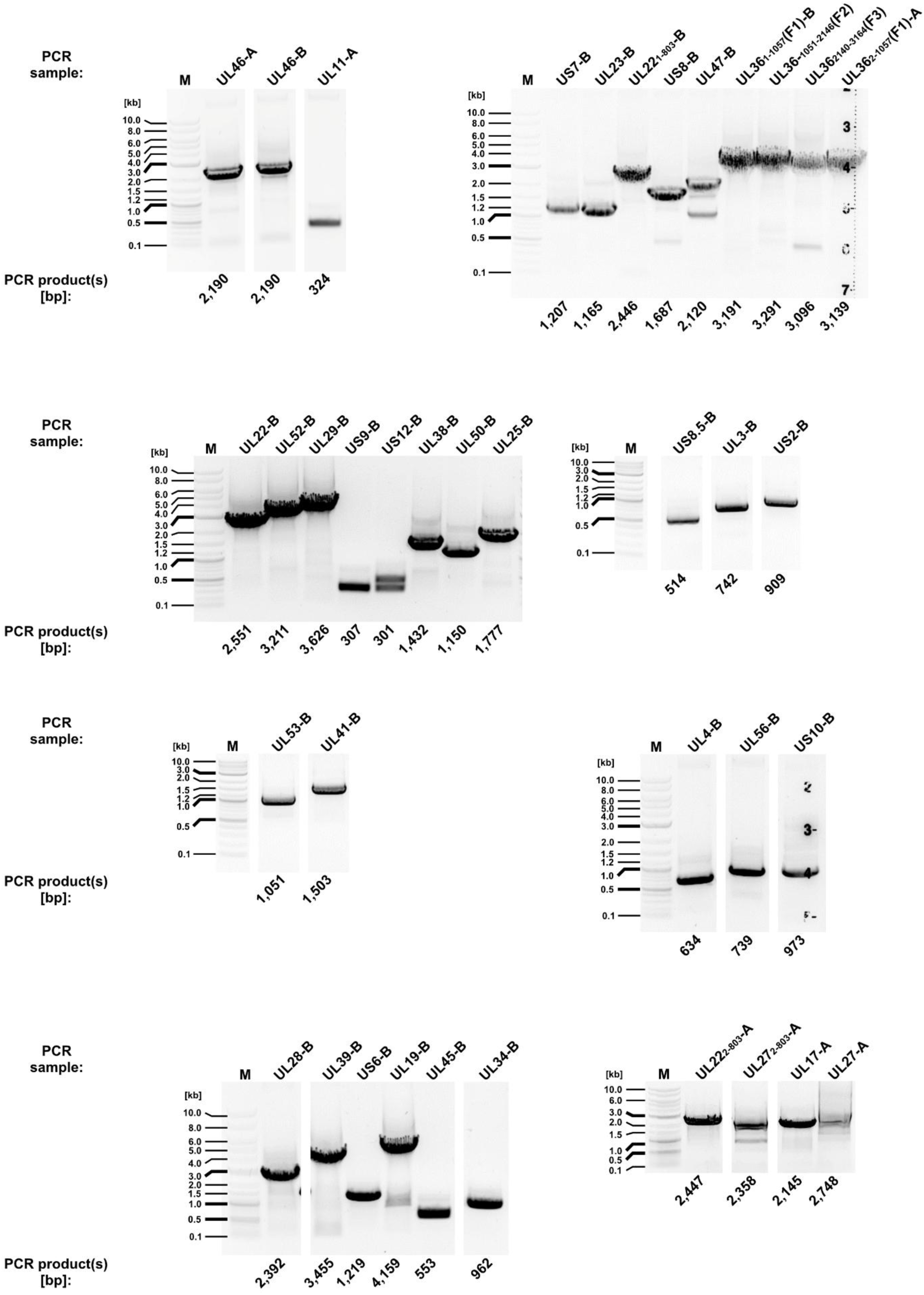
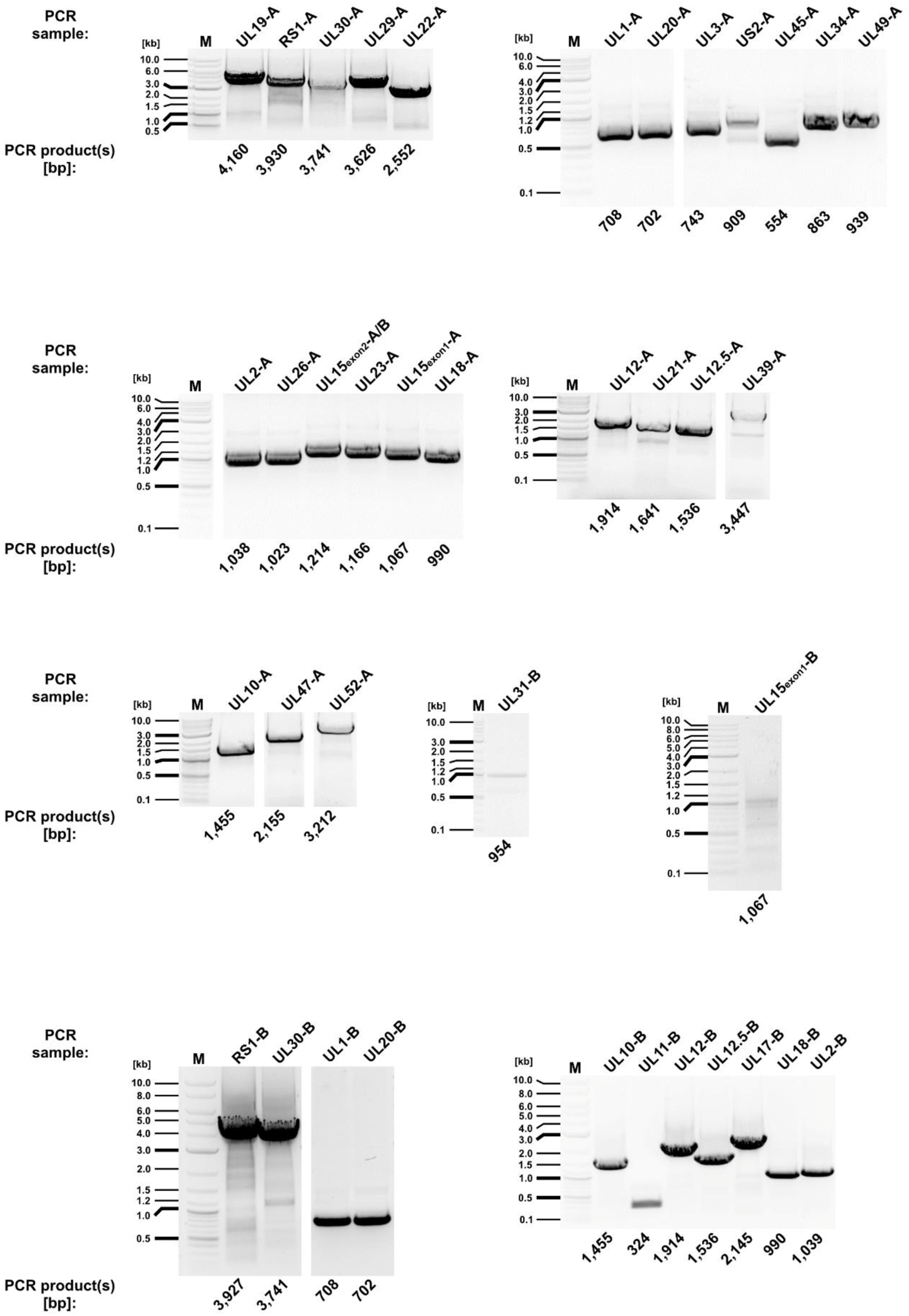
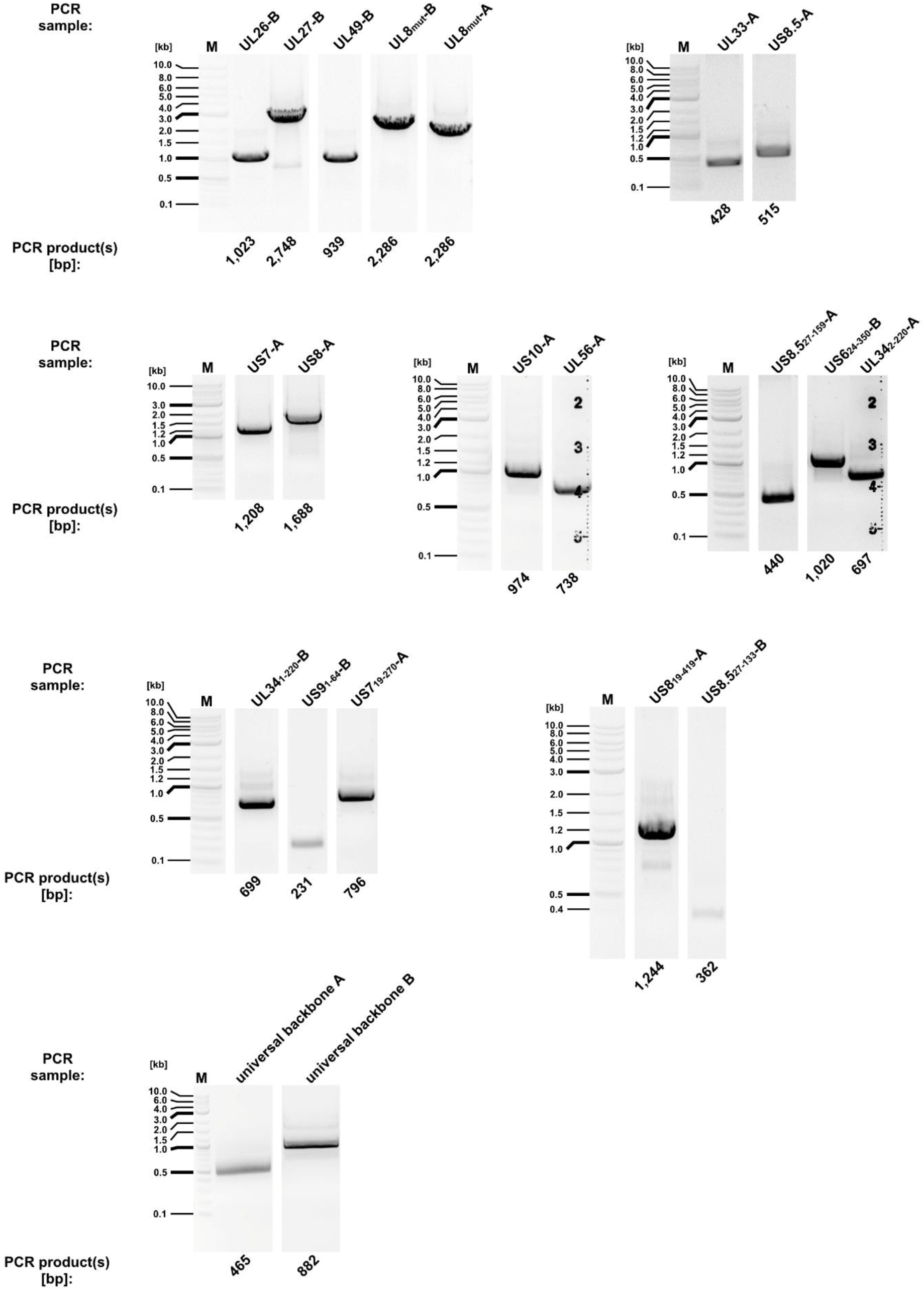
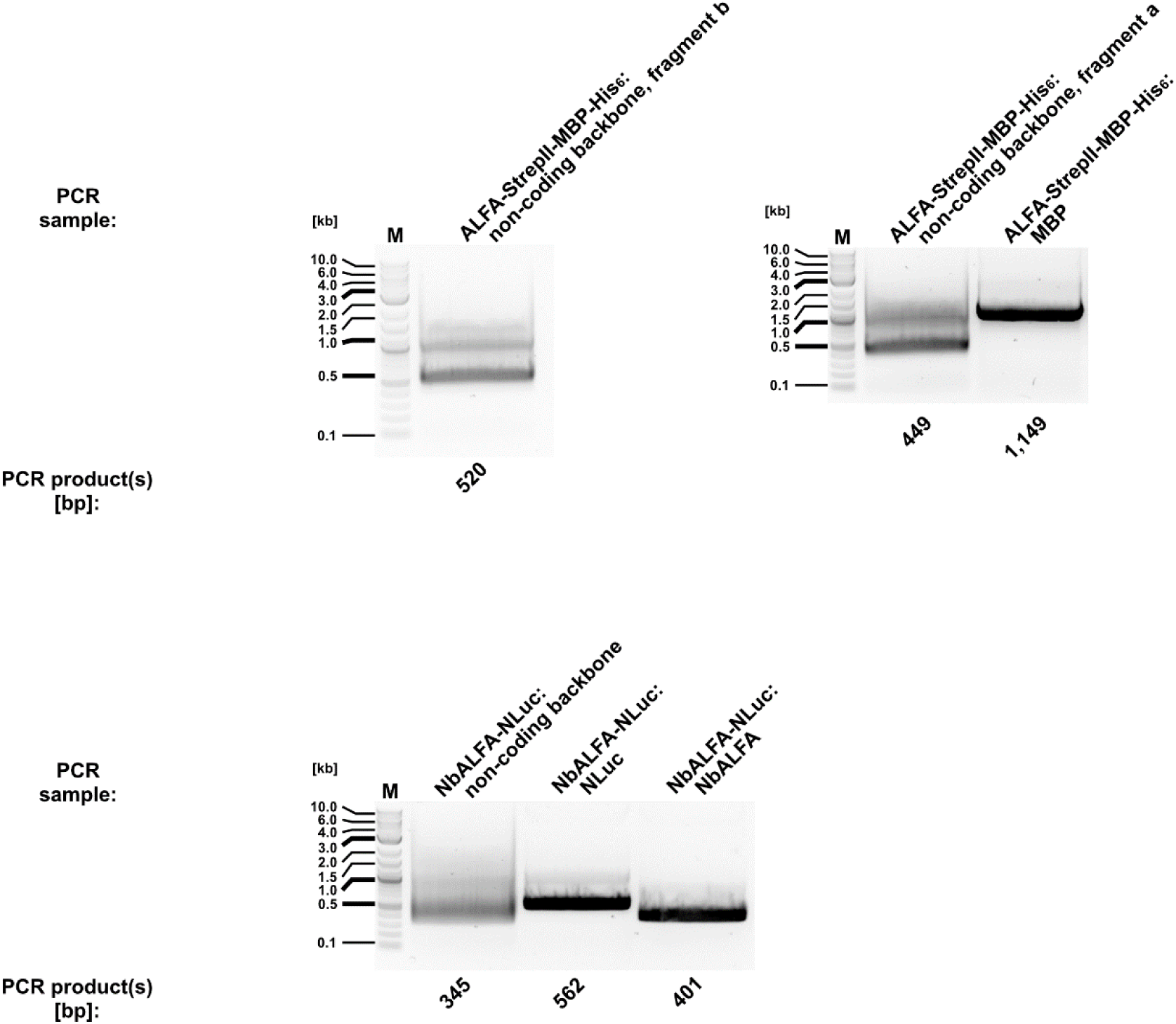
Electrophoretic validation of HSV-1 gene PCR products. Amplicons of the indicated HSV-1 CDSs were resolved by 0.5% or 1.0% (w/v) agarose gel electrophoresis to confirm expected molecular weights. Individual gene fragment names are labeled above the corresponding lanes, with their predicted fragment sizes (in base pairs) indicated below. A representative DNA ladder (M) was used as a molecular weight standard to ensure accurate size estimation.

**Supplemental Figure 7:**
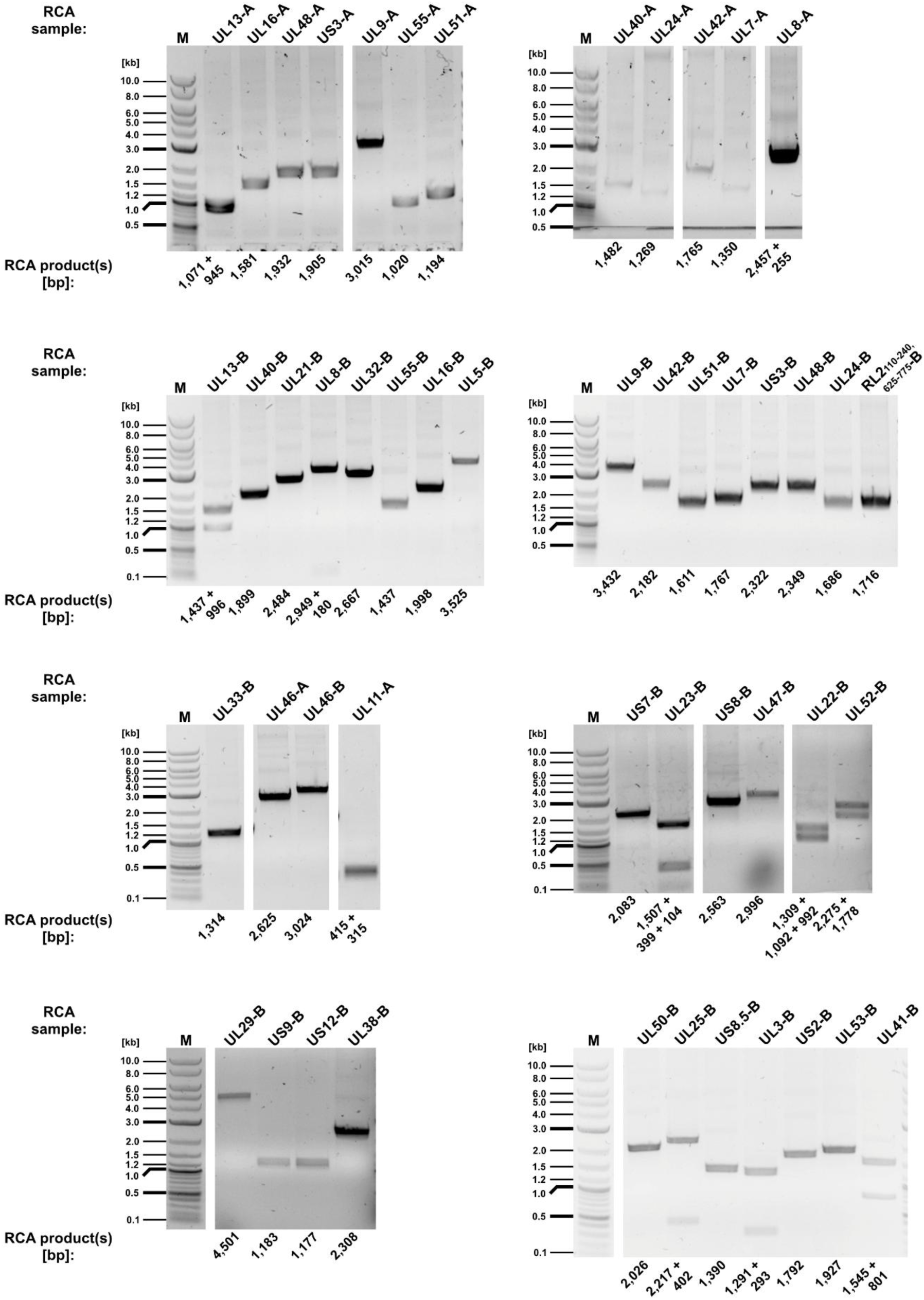
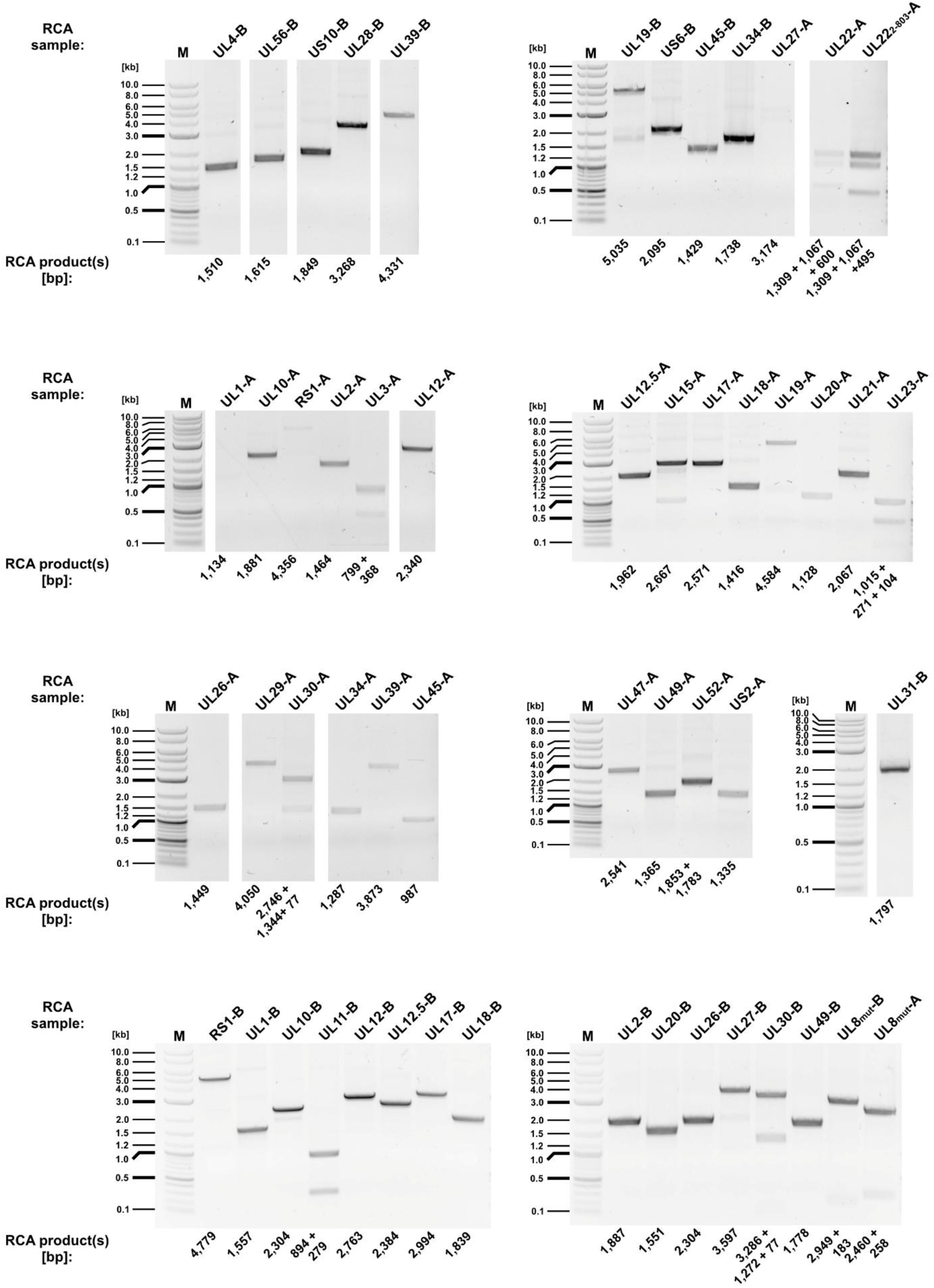
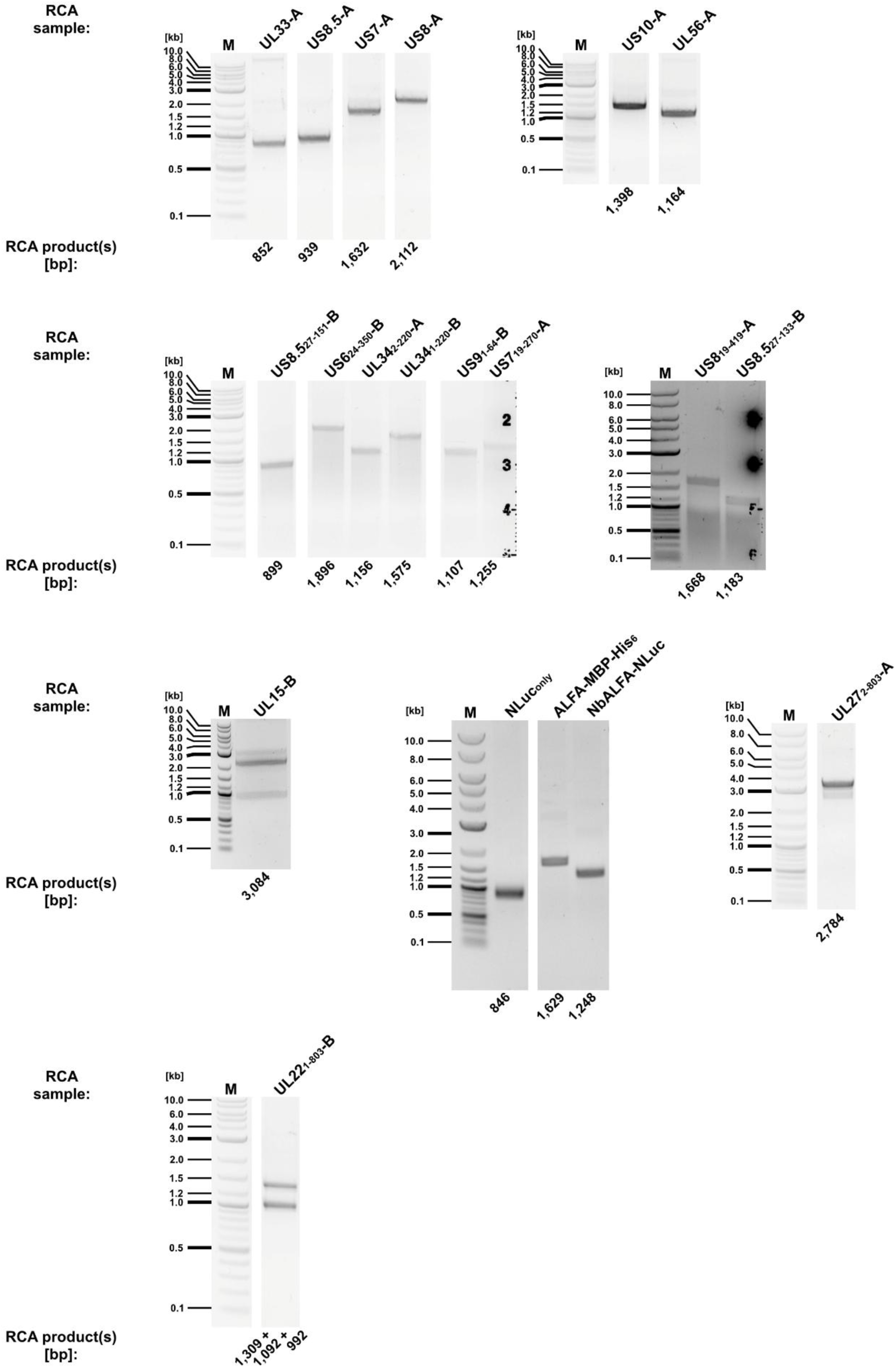
Restriction digestion analysis of RCA products. One microliter of each RCA product was digested with EcoRV and resolved by 0.5% or 1.0% (w/v) agarose gel electrophoresis to confirm expected molecular weights. Individual gene fragment names are labeled above the corresponding lanes, with their predicted digestion product sizes (in base pairs) indicated below. A representative DNA ladder (M) was used as a molecular weight standard to ensure accurate size estimation.

### Cell-Free Protein Expression

Recombinant proteins for the LUCIA assay were synthesized using the myTXTL Pro Cell-Free Expression Kit (Arbor Biosciences, 540300) according to the manufacturer’s protocol. To maintain optimal enzymatic activity, all proteins were expressed immediately prior to use. Briefly, an equivalent of 60 fmol of unit-length RCA template was incorporated into the *E. coli* cell-free master mix, achieving a final reaction volume of 12 μL per sample. Protein synthesis was performed by incubation at 27°C for 18 hours. Following expression, reaction mixtures were maintained on ice for immediate application in downstream assays.

### Determination of His-tagged A Protein Binding Kinetics to Nickel-Coated Plates

To determine the binding kinetics of His-tagged proteins to Pierce Nickel-Coated 96-well plates (Thermo Scientific, 15242), a standard curve was established using conventionally purified ALFA-MBP-His_6_. Increasing concentrations of the purified protein were diluted in PBS to a final volume of 200 μL in protein low binding tubes (Sarstedt, 72.706.600), loaded onto the plates, and incubated for 1 hour at 4°C with orbital shaking at 500 rpm. Following incubation, wells were washed three times with 300 μL PBS-T (PBS containing 0.05% Tween-20). The plates were subsequently incubated with an HRP-conjugated anti-ALFA nanobody (1:5,000 in PBS-T; NanoTag Biotechnologies, N1505-HRP) for 1 hour at room temperature with shaking at 500 rpm. After three additional washes with 300 μL PBS-T, 200 μL of SuperSignal ELISA Pico Chemiluminescent Substrate (Thermo Scientific, 37069) was added to each well. RLUs were measured using a Tecan Infinite 200 PRO M Plex microplate reader precisely 5 minutes after substrate addition. The standard curve was generated using GraphPad Prism 10 software via linear regression, yielding the following equation: Y=2,353,435X+755,748 (Supplemental Fig. 8A and B).

**Supplemental Figure 8:**
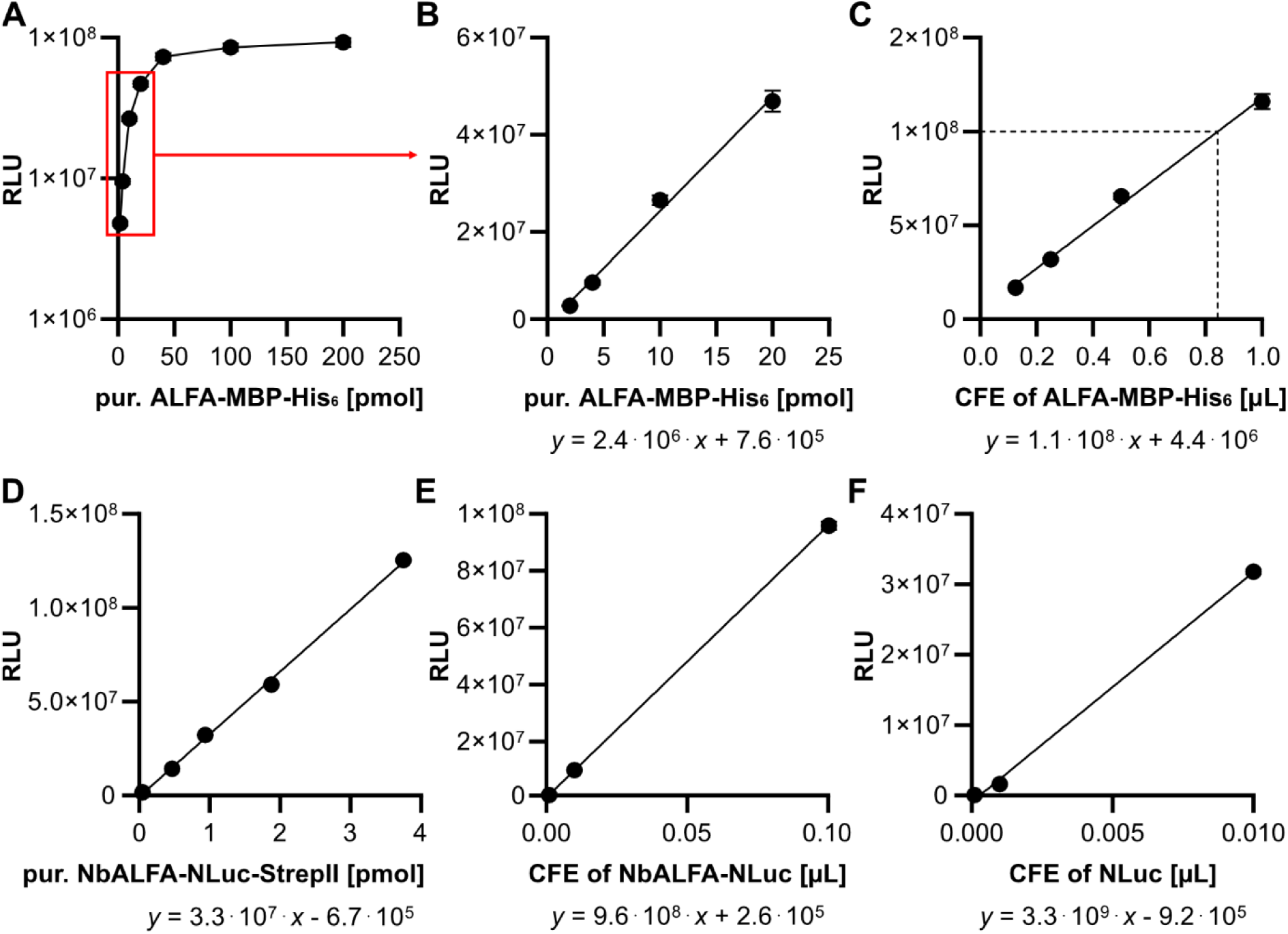
Quantification of proteins produced by cell-free expression. **A**, Binding curve of conventionally purified ALFA-MBP-His_6_ protein to a nickel-coated 96-well plate, shown as RLU as a function of protein input. **B**, Enlarged view of the linear range highlighted in (**A**). **C**, Representative binding curve of a CFE-expressed A protein (ALFA-MBP-His_6_), shown as RLU as a function of CFE lysate input. The vertical dashed line indicates the CFE input volume selected for downstream experiments (equivalent to ∼1×10^8^ RLU), exceeding the estimated binding capacity of a single well (∼10 pmol) as determined from the calibration in (**A**) and (**B**). **D**, Binding curve of conventionally purified ALFA-MBP-His_6_ protein to a nickel-coated 96-well plate, shown as RLU as a function of protein input. **E** and **F**, Representative binding curve of CFE-expressed B proteins NbALFA-NLuc (**E**) and NLuc (**F**), shown as RLU as a function of CFE lysate input.

### Quantification of Cell-Free-Expressed His-tagged A proteins

To quantify the molar concentration of His_6_-tagged A proteins generated via CFE, serial dilutions of fresh, ice-cold translation products (1.0, 0.5, 0.25, and 0.125 μL) were prepared in PBS to a final volume of 200 μL in protein low binding tubes (Sarstedt, 72.706.600). These aliquots were loaded onto Pierce Nickel-Coated 96-well plates (Thermo Fisher Scientific, 15242) and immobilized by incubation for 1 hour at 4°C with orbital shaking at 500 rpm. Following immobilization, wells were washed, stained, and measured as described above. The absolute molar quantities of the His_6_-tagged A proteins within the CFE lysates were subsequently determined by interpolating the RLU values against the previously established linear regression standard curve (Supplemental Fig. 8C). Using these data, the CFE volume of the respective A protein equivalent to ∼1 × 10⁸ RLU was calculated and used in downstream LUCIA experiments.

### Quantification of conventionally purified tagged NLuc protein

To establish the relationship between molar quantity and bioluminescent signal, a standard curve was generated using conventionally purified NbALFA-NLuc-StrepII. A starting amount of 3.75 pmol of protein was serially diluted (1:2) in PBS to a final volume of 100 μL per sample in protein low binding tubes (Sarstedt, 72.706.600). The assay was performed in white 96-well OptiPlates (PerkinElmer, 6055290). Initially, 1 μL Nano-Glo substrate diluted in 10 µL Nano-Glo reaction buffer (Nano-Glo Luciferase Assay System, Promega, N1110) was added to each well, followed by the addition of the protein dilutions. RLUs were measured using a Tecan Infinite 200 PRO M Plex microplate reader precisely 5 minutes after protein addition. The resulting standard curve was modeled in GraphPad Prism 10 using the linear equation: Y=33,326,123X-668,626 (Supplemental Fig. 8D).

### Quantification of Cell-Free-Expressed NLuc-tagged B proteins

For the quantification of NLuc-fused B proteins expressed via CFE, fresh ice-cold translation products (0.01, 0.001, or 0.0001 μL) were diluted in PBS to final volumes of 100 μL. These aliquots were added to plates containing 1 μL Nano-Glo substrate, diluted in 10 µL Nano-Glo reaction buffer. RLU values were recorded precisely 5 minutes after protein addition using a Tecan Infinite 200 PRO M Plex microplate reader. The absolute molar amounts of the CFE-expressed B proteins were subsequently determined by interpolation from the linear regression standard curve (Supplemental Fig. 8E and F). Using these data, the CFE volume of the respective B protein equivalent to ∼4 × 10⁸ RLU was calculated and used in downstream LUCIA experiments to ensure sufficient B protein to saturate available protein A binding sites.

### Quantification of Non-specific Binding

To determine the baseline of non-specific binding for NLuc-fused proteins, all HSV-1 viral proteins meeting the high-confidence PPI criteria (ipTM > 0.6, excluding UL36) were synthesized as NLuc-fused B proteins via CFE. The non-specific background signals for these proteins were quantified as follows (Supplemental Table 2 Sheet 1): Pierce Nickel-Coated 96-well plates were initially blocked with 42 pmol of purified mCherry-His_6_ protein for 1 hour at 4°C with orbital shaking at 500 rpm. Following blocking, the plates were washed three times with 300 μL PBS-T. A standardized quantity of each NLuc-fused B protein, normalized to 4×10^8^ RLU (∼12 pmol), was diluted in PBS-T to a final volume of 200 μL and loaded onto the pre-blocked plates. Samples were incubated for 1 hour at 4°C with orbital shaking at 500 rpm to allow for potential non-specific adherence. After three subsequent washes with 300 μL PBS-T, 1 μL Nano-Glo substrate diluted in 200 µL Nano-Glo reaction buffer was added to each well. Non-specific RLU was recorded using a Tecan Infinite 200 PRO M Plex microplate reader precisely 5 minutes after substrate addition.

### Direct PPI validation with LUminescent Cell-free Interaction Assay (LUCIA)

To validate high-confidence HSV-1 PPI candidates (defined as ipTM > 0.6), the interaction partner exhibiting the lowest non-specific plate binding was selected for expression as the NLuc-fused B protein. All CFE products were freshly synthesized and quantified immediately prior to the assay to ensure consistent input. The LUCIA procedure was initiated by immobilizing 1×10^8^ RLU (approximately 42 pmol) of His-tagged A proteins, diluted in 200 µL of PBS in protein low binding tubes (Sarstedt, 72.706.600), onto Pierce Nickel-Coated 96-well plates. Incubation was performed for 1 hour at 4°C with orbital shaking at 500 rpm. To provide a rigorous baseline for non-specific binding, 42 pmol of purified mCherry-His_6_ was processed in parallel under identical conditions. Following immobilization, wells were washed three times with 300 µL of PBS-T. For the interaction phase, 4×10^8^ RLU (approximately 12 pmol) of NLuc-fused B proteins were diluted in 200 µL of PBS-T in protein low binding tubes (Sarstedt, 72.706.600), added to the respective wells, and incubated for 1 hour at 4°C with shaking at 500 rpm. After three subsequent washes with 300 µL of PBS-T, 1 μL Nano-Glo substrate diluted in 200 µL Nano-Glo reaction buffer was added to each well. RLU was measured using a Tecan Infinite 200 PRO M Plex microplate reader precisely 5 minutes after substrate addition. Binding affinity was quantified by calculating the fold change (FC) of the experimental signal relative to the mCherry-His_6_ background control.

### Cell Transfection and Protein Pull-down

For mammalian protein expression, HEK 293XT cells were seeded in 10 cm dishes and transfected with 8 µg of total plasmid DNA using PEI. At 24 hpt, cells were washed twice with ice-cold PBS and incubated on ice for 30 minutes in 1 mL of NP-40 lysis buffer [50 mM Tris-HCl (pH 7.5), 150 mM NaCl, 1% Nonidet P-40 (v/v)] supplemented with cOmplete Protease Inhibitor Cocktail (Roche, 11836170001). The resulting lysates were cleared of cellular debris by centrifugation at 10,000 xg for 20 minutes at 4°C. His-tagged proteins were isolated from the supernatant using His-Tag Affinity MagBeads (Cube Biotech, 31205) according to the manufacturer’s instructions. Samples were resolved by SDS-PAGE and analyzed via Western blot. The following antibodies were utilized at a 1:5,000 dilution: Alexa Fluor 647-conjugated anti-6xHis (BioLegend, 362611), anti-ALFA-tag-HRP (NanoTag, N1505), and anti-α-tubulin (Calbiochem, CP06) and Alexa Fluor 647-conjugated goat anti-mouse IgG (H+L) (Invitrogen; A21236) as a loading control. Chemiluminescent signals were captured using a FUSION FX7 EDGE 18.11 imaging system.

### Immunofluorescence staining

Vero B4 cells were seeded in 24-well plates (Sarstedt, 83.3922) at a density of 1×10^5^ cells per well. After 24 h, cells were transfected with 400 ng of BAC DNA per well using Lipofectamine 2000 (Invitrogen, 11668027) according to the manufacturer’s protocol. Cells were fixed at 24 h intervals up to 96 hpt using 5% paraformaldehyde (w/v) (Electron Microscopy Sciences, 15713-S) in PBS for 15 min at 37°C. Following fixation, cells were permeabilized with 0.1% (v/v) Triton X-100 (Sigma-Aldrich, T8787-50ML) in PBS for 20 min at RT and subsequently blocked with 3% (w/v) bovine serum albumin (BSA; Millipore, 81-053-3) in PBS. For viral ICP4 protein detection, cells were incubated with a primary antibody against ICP4 (1:500; Santa Cruz Biotechnology, sc-56986) in PBS supplemented with 1% (w/v) BSA for 1 h at RT. This was followed by incubation with an Alexa Fluor 488-conjugated secondary antibody (1:500; Invitrogen, A11001) for 1 h at RT. Nuclei were counterstained with Hoechst 33342 (1:1,000) for 20 min at RT. Samples were maintained in PBS at 4°C until imaging. Fluorescence images were captured using a Leica DM IL LED inverted widefield microscope equipped with a Leica DMC 4500 camera.

### Data Analysis and Software

Analysis of AF2 predictions was performed with a previously described python script^9^. A one-sided *t*-test for the LUCIA ratios was performed with the python scipy package. A significance threshold of *P* value < 0.05 was used. Structural data were visualized and analyzed using UCSF ChimeraX v1.9 (https://www.cgl.ucsf.edu/chimerax). FoldX 4 was used to design the HSV-1 UL8 mutant.

### Use of large language models

Large Language Models (Claude, Gemini 3.0 Pro, and GPT 5.2) were employed during the drafting process to optimize editorial content, including grammatical correction and stylistic flow. The authors manually verified all AI-assisted text revisions and assume full responsibility for the scientific integrity and content of the published article.

## Supplemental Tables

### Supplemental Table 1. In silico scores and evaluation of quality

Sheet 1) AlphaFold2 ipTM scores of the interactome predictions of HSV-1, HCMV, and KSHV.

Sheet 2) Receiver operating characteristic (ROC) analysis of ipTM on E. coli dimer database. The ipTM of the positive and negative populations are provided as well as the Youden index to determine the optimal ipTM threshold and the area under the curve (AUC) statistics.

Sheet 3) Evaluation of actifpTM and ipSAE to identify LUCIA positive protein pairs.

Sheet 4) Recall calculations for ipTM and LUCIA using PDB structures as the true positives.

Sheet 5) AlphaFold2 ipTM scores of the ectodomain-only transmembrane proteins tested in LUCIA.

Sheet 6) FoldX analysis and AlphaFold2 ipTM scores for the mutations in HSV-1 UL8.

### Supplemental Table 2. LUCIA bioluminescence data and primer sequences for HSV-1 PPI validation

Sheet 1) Raw bioluminescence values of unspecific background binding of B proteins.

Sheet 2) LUCIA raw bioluminescence values of high-confidence HSV-1 PPIs including background controls, used for recall calculations.

Sheet 3) LUCIA raw bioluminescence values of full length transmembrane proteins.

Sheet 4) List of primers used for BAC mutagenesis.

Sheet 5) List of primers used for amplification of HSV-1 CDSs.

## Data Availability

The HSV-1, HCMV, and KSHV protein structure predictions have been deposited under DOI doi.org/10.5281/zenodo.12517508 at https://zenodo.org/records/12517508. The additional AF2 models, *i.e. E. coli* ROC, UL8 mutants, ectodomain-only transmembrane proteins, and rerun of PDB dimers, and supporting LUCIA documents, *i.e.* cloning validation, associated plasmid maps, and DNA sequences, have been deposited in the Zenodo database under DOI 10.5281/zenodo.19328587 at https://zenodo.org/records/19328587. Furthermore, an interactive web interface is available at https://www.herpesfolds.org/herpesppis.

## Acknowledgements

We acknowledge the Maxwell computational resources operated at Deutsches Elektronen-Synchrotron (DESY), Hamburg, Germany, and the resources provided by the Medizinische Hochschule Hannover (MHH) Information Technology (MIT) High Performance Computing (HPC) team.

Work in the Bosse lab was funded by the Deutsche Forschungsgemeinschaft (DFG, German Research Foundation) under Germany’s Excellence Strategy EXC 2155 project no. 390874280, the DFG-funded RTG 2771 Humans and Microbes, project number 453548970, the DFG-funded RTG 2887 VISION, project number 49735088, DFG-funded CRC 1648 Emerging Viruses project number SFB 1648/1 2024-512741711, by the Wellcome Trust through a Collaborative Award (209250/Z/17/Z), and the Leibniz ScienceCampus InterACt, funded by the BWFGB Hamburg and the Leibniz Association (W75/2022) InterACt and “Hamburg- X Infektionsforschung”. Moreover, the Bosse lab is funded through the DFG Research Unit FOR5200 DEEP-DV (443644894) project BO 4158/5-1 and BO 4158/5-2, DFG Research Unit FOR5898 AdBHealth (548065690) project BO 4158/9-1 and the German Center for Infection Research (DZIF) grants IICH TTU07.918, TTU 07.861 and 07.863.

## Author Contributions

**Tianyu Zhang**: Conceptualization, Data Curation, Formal Analysis, Investigation, Methodology, Validation, Visualization, Writing - original draft, Writing - review & editing; **Julian Kraft**: Conceptualization, Data Curation, Formal Analysis, Investigation, Methodology, Validation, Visualization, Writing - original draft, Writing - review & editing; **Timothy K. Soh**: Conceptualization, Data Curation, Formal Analysis, Investigation, Methodology, Software, Validation, Visualization, Writing - original draft, Writing - review & editing; **Malte Kansy**: Data Curation, Resources, Software; **Ingvar Jonsson**: Investigation, Methodology, Software; **Jens B. Bosse**: Conceptualization, Funding Acquisition, Methodology, Project Administration, Supervision, Validation, Writing - original draft, Writing - review & editing

## References

1. Nooren, I. M. A. & Thornton, J. M. Diversity of protein-protein interactions. EMBO J. 22, 3486–3492 (2003).

2. Krogan, N. J. et al. Global landscape of protein complexes in the yeast Saccharomyces cerevisiae. Nature 440, 637–643 (2006).

3. Porta-Pardo, E., Ruiz-Serra, V., Valentini, S. & Valencia, A. The structural coverage of the human proteome before and after AlphaFold. PLoS Comput. Biol. 18, e1009818 (2022).

4. Bayly-Jones, C. & Whisstock, J. C. Mining folded proteomes in the era of accurate structure prediction. PLoS Comput. Biol. 18, e1009930 (2022).

5. Jumper, J. et al. Highly accurate protein structure prediction with AlphaFold. Nature 596, 583–589 (2021).

6. Abramson, J. et al. Accurate structure prediction of biomolecular interactions with AlphaFold 3. Nature 630, 493–500 (2024).

7. Baek, M. et al. Accurate prediction of protein structures and interactions using a three-track neural network. Science 373, 871–876 (2021).

8. Tunyasuvunakool, K. et al. Highly accurate protein structure prediction for the human proteome. Nature 596, 590–596 (2021).

9. Soh, T. K. et al. A proteome-wide structural systems approach reveals insights into protein families of all human herpesviruses. Nat. Commun. 15, 10230 (2024).

10. Bordin, N. et al. Novel machine learning approaches revolutionize protein knowledge. Trends Biochem. Sci. 48, 345–359 (2023).

11. Terwilliger, T. C. et al. AlphaFold predictions are valuable hypotheses and accelerate but do not replace experimental structure determination. Nat. Methods 21, 110–116 (2024).

12. Zhang, J. et al. Predicting protein-protein interactions in the human proteome. Science 390, eadt1630 (2025).

13. Schmid, E. W. & Walter, J. C. Predictomes, a classifier-curated database of AlphaFold-modeled protein-protein interactions. Mol. Cell 85, 1216–1232.e5 (2025).

14. Farooq, A. V. & Shukla, D. Herpes simplex epithelial and stromal keratitis: an epidemiologic update. Surv. Ophthalmol. 57, 448–462 (2012).

15. Korver, A. M. H. et al. Congenital hearing loss. Nat. Rev. Dis. Primer 3, 16094 (2017).

16. Ganem, D. KSHV and the pathogenesis of Kaposi sarcoma: listening to human biology and medicine. J. Clin. Invest. 120, 939–949 (2010).

17. Davison, A. J. Herpesvirus systematics. Vet. Microbiol. 143, 52–69 (2010).

18. Shah, P. S. et al. Systems Biology of Virus-Host Protein Interactions: From Hypothesis Generation to Mechanisms of Replication and Pathogenesis. Annu. Rev. Virol. 9, 397–415 (2022).

19. Dogrammatzis, C., Waisner, H. & Kalamvoki, M. ‘Non-Essential’ Proteins of HSV-1 with Essential Roles In Vivo: A Comprehensive Review. Viruses 13, 17 (2020).

20. Nobre, L. V. et al. Human cytomegalovirus interactome analysis identifies degradation hubs, domain associations and viral protein functions. eLife 8, e49894 (2019).

21. Bogdanow, B. et al. Structural host-virus interactome profiling of intact infected cells. Nat. Commun. 16, 6713 (2025).

22. Licata, L. et al. MINT, the molecular interaction database: 2012 update. Nucleic Acids Res. 40, D857–861 (2012).

23. Del Toro, N. et al. The IntAct database: efficient access to fine-grained molecular interaction data. Nucleic Acids Res. 50, D648–D653 (2022).

24. Hernández Durán, A., Grünewald, K. & Topf, M. Conserved Central Intraviral Protein Interactome of the Herpesviridae Family. mSystems 4, e00295–19 (2019).

25. EMBL. Confidence scores in AlphaFold-Multimer. Confidence scores in AlphaFold-Multimer https://www.ebi.ac.uk/training/online/courses/alphafold/inputs-and-outputs/evaluating-alphafolds-predicted-structures-using-confidence-scores/confidence-scores-in-alphafold-multimer/ (2026).

26. Yin, R., Feng, B. Y., Varshney, A. & Pierce, B. G. Benchmarking AlphaFold for protein complex modeling reveals accuracy determinants. Protein Sci. Publ. Protein Soc. 31, e4379 (2022).

27. O’Reilly, F. J. et al. Protein complexes in cells by AI-assisted structural proteomics. Mol. Syst. Biol. 19, e11544 (2023).

28. Götzke, H. et al. The ALFA-tag is a highly versatile tool for nanobody-based bioscience applications. Nat. Commun. 10, 4403 (2019).

29. Rajagopala, S. V. et al. The binary protein-protein interaction landscape of Escherichia coli. Nat. Biotechnol. 32, 285–290 (2014).

30. Varga, J. K., Ovchinnikov, S. & Schueler-Furman, O. actifpTM: a refined confidence metric of AlphaFold2 predictions involving flexible regions. Bioinformatics 41, btaf107 (2025).

31. Dunbrack, R. L. Rēs ipSAE loquunt: What’s wrong with AlphaFold’s ipTM score and how to fix it. BioRxiv Prepr. Serv. Biol. 2025.02.10.637595 (2025) doi:10.1101/2025.02.10.637595.

32. Dai, X. & Zhou, Z. H. Structure of the herpes simplex virus 1 capsid with associated tegument protein complexes. Science 360, eaao7298 (2018).

33. Mapelli, M., Panjikar, S. & Tucker, P. A. The crystal structure of the herpes simplex virus 1 ssDNA-binding protein suggests the structural basis for flexible, cooperative single-stranded DNA binding. J. Biol. Chem. 280, 2990–2997 (2005).

34. Bigalke, J. M., Heuser, T., Nicastro, D. & Heldwein, E. E. Membrane deformation and scission by the HSV-1 nuclear egress complex. Nat. Commun. 5, 4131 (2014).

35. Boehmer, P. E. & Nimonkar, A. V. Herpes virus replication. IUBMB Life 55, 13–22 (2003).

36. Yu, Z. et al. Mechanisms of HSV-1 helicase-primase inhibition and replication fork complex assembly. Cell 189, 478–494.e18 (2026).

37. Bermek, O., Willcox, S. & Griffith, J. D. DNA replication catalyzed by herpes simplex virus type 1 proteins reveals trombone loops at the fork. J. Biol. Chem. 290, 2539–2545 (2015).

38. Meng, Y. et al. Protein structure prediction via deep learning: an in-depth review. Front. Pharmacol. 16, 1498662 (2025).

39. Kawaguchi, S. et al. In silico screening by AlphaFold2 program revealed the potential binding partners of nuage-localizing proteins and piRNA-related proteins. eLife 13, RP101967 (2025).

40. Homma, F., Huang, J. & van der Hoorn, R. A. L. AlphaFold-Multimer predicts cross-kingdom interactions at the plant-pathogen interface. Nat. Commun. 14, 6040 (2023).

41. Abulude, I. J. et al. Using AlphaFold-Multimer to study novel protein-protein interactions of predation essential hypothetical proteins in Bdellovibrio. Front. Bioinforma. 5, 1566486 (2025).

42. Bogdanow, B. et al. Spatially resolved protein map of intact human cytomegalovirus virions. Nat. Microbiol. 8, 1732–1747 (2023).

43. Leroy, B., Gillet, L., Vanderplasschen, A. & Wattiez, R. Structural Proteomics of Herpesviruses. Viruses 8, 50 (2016).

44. Bryant, P. et al. Predicting the structure of large protein complexes using AlphaFold and Monte Carlo tree search. Nat. Commun. 13, 6028 (2022).

45. Apweiler, R. et al. UniProt: the Universal Protein knowledgebase. Nucleic Acids Res. 32, D115–119 (2004).

46. Steinegger, M. & Söding, J. MMseqs2 enables sensitive protein sequence searching for the analysis of massive data sets. Nat. Biotechnol. 35, 1026–1028 (2017).

47. van Kempen, M. et al. Fast and accurate protein structure search with Foldseek. Nat. Biotechnol. 42, 243–246 (2024).

48. Skwarczynska, M. & Ottmann, C. Protein-protein interactions as drug targets. Future Med. Chem. 7, 2195–2219 (2015).

49. Higueruelo, A. P., Jubb, H. & Blundell, T. L. Protein-protein interactions as druggable targets: recent technological advances. Curr. Opin. Pharmacol. 13, 791–796 (2013).

50. Nada, H. et al. New insights into protein-protein interaction modulators in drug discovery and therapeutic advance. Signal Transduct. Target. Ther. 9, 341 (2024).

51. Segal-Maurer, S. et al. Capsid Inhibition with Lenacapavir in Multidrug-Resistant HIV-1 Infection. N. Engl. J. Med. 386, 1793–1803 (2022).

52. Sandbaumhüter, M. et al. Cytosolic herpes simplex virus capsids not only require binding inner tegument protein pUL36 but also pUL37 for active transport prior to secondary envelopment: The role of pUL36 and pUL37 for HSV1 capsid transport. Cell. Microbiol. 15, 248–269 (2013).

53. Bosse, J. B. et al. Remodeling nuclear architecture allows efficient transport of herpesvirus capsids by diffusion. Proc. Natl. Acad. Sci. 112, (2015).

54. Tischer, B. K., Smith, G. A. & Osterrieder, N. En passant mutagenesis: a two step markerless red recombination system. Methods Mol. Biol. 634, 421–430 (2010).

55. Mirdita, M. et al. ColabFold: making protein folding accessible to all. Nat. Methods 19, 679–682 (2022).

